# Super-immunity by broadly protective nanobodies to sarbecoviruses

**DOI:** 10.1101/2021.12.26.474192

**Authors:** Yufei Xiang, Wei Huang, Hejun Liu, Zhe Sang, Sham Nambulli, Jérôme Tubiana, Kevin L Williams, W Paul Duprex, Dina Schneidman-Duhovny, Ian A. Wilson, Derek J. Taylor, Yi Shi

**Affiliations:** Department of Cell Biology, University of Pittsburgh, Pittsburgh, PA, 15213, USA; Department of Pharmacology, Case Western Reserve University, Cleveland, OH 44106, USA; Department of Integrative Structural and Computational Biology, The Scripps Research Institute, La Jolla, CA, 92037, USA; The University of Pittsburgh and Carnegie Mellon University Program for Computational Biology, Pittsburgh, PA, 15213, USA; Center for Vaccine Research, University of Pittsburgh, Pittsburgh, PA, 15213, USA; Department of Microbiology and Molecular Genetics, University of Pittsburgh, Pittsburgh, PA, 15213, USA; School of Computer Science and Engineering, The Hebrew University of Jerusalem, Israel; Blavatnik School of Computer Science, Tel Aviv University, Israel; Skaggs Institute for Chemical Biology, The Scripps Research Institute, La Jolla, CA, 92037, USA; Department of Biochemistry, Case Western Reserve University, Cleveland, OH 44106, USA

## Abstract

Vaccine boosters and infection can facilitate the development of SARS-CoV-2 antibodies with improved potency and breadth. Here, we observed super-immunity in a camelid extensively immunized with the SARS-CoV-2 receptor-binding domain (RBD). We rapidly isolated a large repertoire of specific ultrahigh-affinity nanobodies that bind strongly to all known sarbecovirus clades using integrative proteomics. These pan-sarbecovirus nanobodies (psNbs) are highly effective against SARS-CoV and SARS-CoV-2 variants including the Omicron, with the best median neutralization potency at single-digit ng/ml. Structural determinations of 13 psNbs with the SARS-CoV-2 spike or RBD revealed five epitope classes, providing insights into the mechanisms and evolution of their broad activities. The highly evolved psNbs target small, flat, and flexible epitopes that contain over 75% of conserved RBD surface residues. Their potencies are strongly and negatively correlated with the distance of the epitopes to the receptor binding sites. A highly potent, inhalable and bispecific psNb (PiN-31) was developed. Our findings inform on the development of broadly protective vaccines and therapeutics.

**One sentence summary:** Successive immunization of SARS-CoV-2 RBD in a camelid enhanced the development of super-immunity and isolation and systematic characterization of a large repertoire of ultrahigh-affinity pan-sarbecovirus single-chain V_H_H antibodies to understand the evolution of this potent and broad immune response.

## MAIN

SARS-related coronaviruses (sarbecoviruses) including SARS-CoV and SARS-CoV-2, are among the most pressing threats to global health. The high genetic diversity, frequent recombination, large natural reservoirs, and proximity to heavily populated areas across continents, underlie the recurrent zoonotic risks of sarbecoviruses and other circulating coronaviruses (*1, 2*). Thus, there is an urgent need to develop broad, effective, and complementary interventions against the currently evolving pandemic as well as future threats. Emerging evidence indicates that B cells isolated from convalescent and infected-then-vaccinated individuals continue to evolve, producing antibodies with increased potency against SARS-CoV-2 antigenic drift (*3–7*). Most antibodies target the spike RBD, which dominates immunogenicity and neutralizing activities of convalescent and vaccinated sera (*8*). Only a small number of Immunoglobulin G (IgG) antibodies against the large sarbecovirus family have been successfully isolated with varied breadth and potency (*9–13*). Moreover, an understanding of the full dynamic repertoire of broadly neutralizing antibodies and mechanisms that shape SARS-CoV-2 super-immunity is still lacking (*3*).

By camelid immunization and proteomics, we recently identified thousands of high-affinity RBD nanobodies (Nbs) that potently neutralize SARS-CoV-2 (*14*). Most ultrapotent Nbs bind the highly variable human ACE2 (hACE2) receptor-binding sites (RBS) and are therefore less effective against evolving variants. Here, we found that, after immune boosters with recombinant RBD_SARS-CoV-2_, serum V_H_H antibodies evolved with substantially improved activities not only against the variants of concern (VOCs) but also to a broad spectrum of sarbecoviruses. To understand the broad serologic activities, we isolated and systematically characterized 100 high-affinity pan-sarbecovirus nanobodies (psNbs) with superior potency and breadth and developed an ultrapotent, bispecific and aerosolizable psNb (PiN-31). Structural determinations of 13 diverse psNbs with the spike or RBD by cryoEM and X-ray crystallography revealed five classes with marked diversity within dominant classes. Our analysis offers insights into the remarkable evolution of serologic responses towards broad activity against sarbecoviruses.

### Identification and characterization of a large repertoire of potent pan-sarbecovirus Nbs

A llama was initially immunized with SARS-CoV-2 RBD-Fc fusion protein and the initial bleed was collected approximately 2 months after priming and three boosts (*14*). We then re-immunized with four additional boosts for 2 months before the booster bleed was collected (see **Methods** in **Supplementary Materials**). The polyclonal V_H_H mixture of the booster bleed showed higher affinity to RBD_SARS-CoV-2_ (ELISA IC_50_s of 43 pM) compared to that of the initial bleed (IC_50_ of 130 pM) (**Fig. 1B**, **fig. S1A**). The V_H_H mixture also maintained excellent neutralization potency (IC_50_s between 0.3-0.6 nM) against the Wuhan-Hu-1 strain as well as Alpha and Lambda VOCs (**fig. S1B**. Notably, compared to the initial V_H_Hs, the neutralization potencies after booster were substantially increased to Beta, Delta and SARS-CoV (IC_50_s of 0.58, 1.90 and 1.65 nM, respectively) corresponding to 6.0, 2.3, and 9.3 fold improvements (**fig. S1B**). Surprisingly, the booster was also associated with strong and broad activities against the full spectrum of sarbecoviruses (**Fig. 1, A and B**). The relative binding affinities (ELISA IC_50_s) for RBDs from clade 1a (RBD_SARS-CoV_), 2 (RBD_RmYN02-CoV_), and 3 (RBD_BM-4831-CoV_) were 0.08, 0.09, and 0.14 nM, respectively, which correspond to 8.5, 7.2, and 8.6 fold improvements over the initial bleed (**fig. S1A**).

**Fig. 1.**
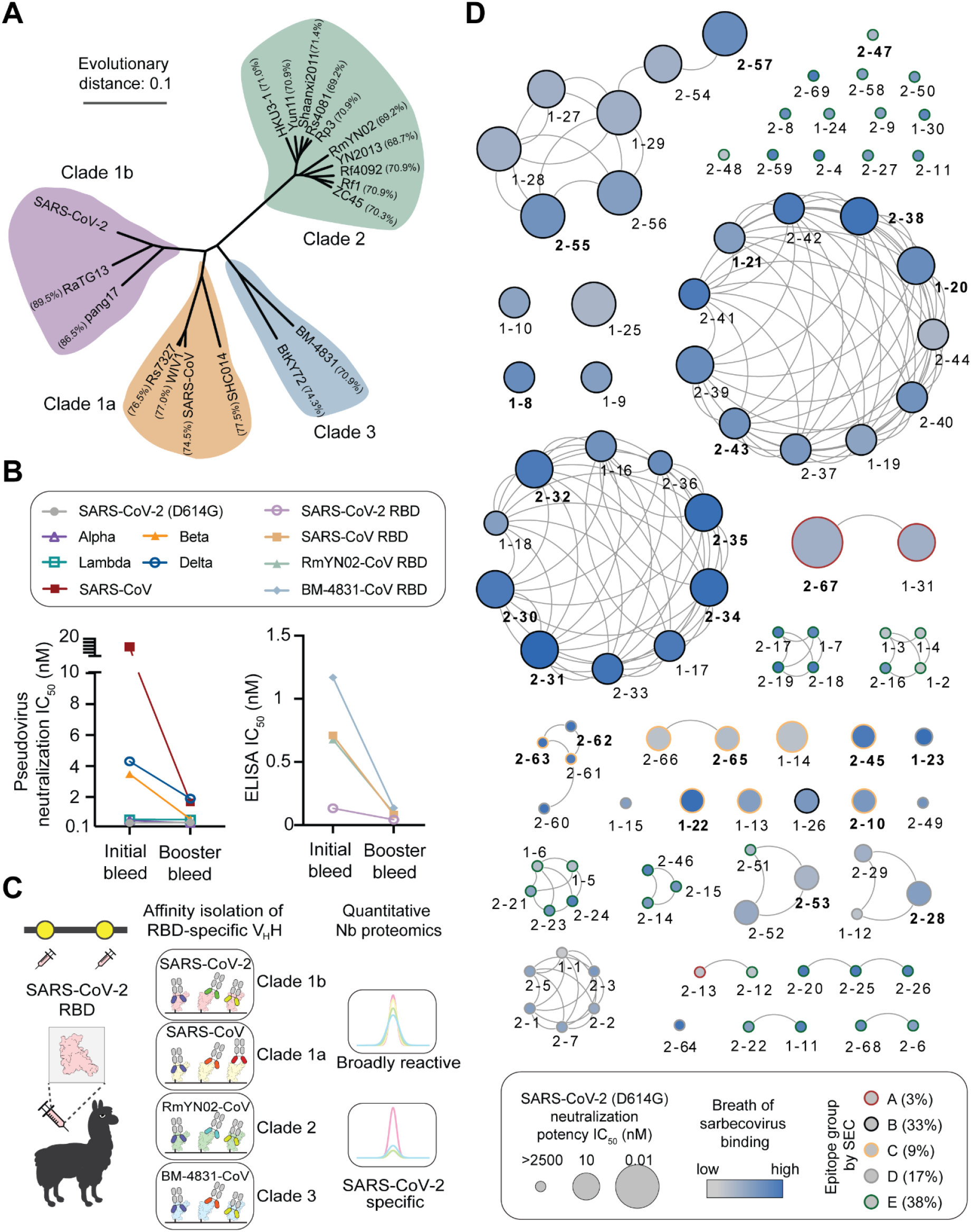
Identification and characterization of psNbs. **(A)** Phylogenetic tree of 19 RBDs from all 4 clades of sarbecoviruses, constructed by the maximum likelihood. **(B)** The neutralization of polyclonal VHHs from two immunization bleeds against pseudovirus of SARS-CoV-2, its variants and SARS-CoV. Their ELISA IC_50_s against 4 RBD clades were also shown. **(C)** Schematics for proteomic identification of psNbs from immunized sera. **(D)** A map summarizing RBD binding and neutralization for 100 high-affinity psNbs. Nbs are represented by dots of various sizes and colors. Two Nbs are connected if their CDR sequence identity is > 70%. Their neutralization potencies against pseudotyped SARS-CoV-2 (D614G) are represented by the size of dots. Breadth of sarbecovirus RBD binding is indicated by the filled gradient color (**table S1**) and the SEC epitope groups (based on benchmark RBD Nbs) are colored on the outer circle (**fig. S3, a and b**). The highest Nb concentration used for binding and neutralization was 8 µM and 2.5 µM, respectively. Bolded psNbs were selected for downstream characterizations. psNbs isolated from the initial and booster bleed were denoted as 1-, and 2- respectively.

Next, we employed quantitative Nb proteomics (*15*) to identify high-affinity psNbs that confer broad-spectrum activity in immunized sera (see **Methods**). This technology rapidly identified hundreds of distinct CDR3 families, from which a fraction of highly divergent Nbs were expressed and characterized. A total of 100 Nbs that interacted strongly with RBD_SARS-CoV-2_ (Clade 1b) were confirmed to cross-react with other sarbecovirus clades (**Fig. 1C, table S1**). Of these, 23%, 35%, and 42% were found to bind two, three, and all four sarbecovirus clades, respectively. Consistent with total polyclonal V_H_H activity, psNbs isolated from the booster show broader pan-sarbecovirus activity than the initial bleed (**fig. S2B**). A substantial fraction (42%) was able to potently neutralize SARS-CoV-2 below 500 nM, with the best IC_50_ of 1.2 ng/ml (77 pM, 2-67), and a median of 0.25 µg/ml (17 nM) (**table S1**). Network analysis revealed that the psNbs are dominated by multiple clusters that span a large spectrum of physicochemical properties including isoelectric point (pI) and hydropathy (**Fig. 1D**, **fig. S2A**). The three largest clusters are each composed of potent neutralizers (with the best IC_50_ of 6.8 ng/ml) that bind strongly to at least three sarbecovirus clades (**table S1**). A smaller cluster (represented by 2-62 and 2-63) showed broad activity, yet only weakly neutralized SARS-CoV-2 *in vitro* (IC_50_s between 40 -132 µg/ml, **table S1, fig. S6**). Other small clusters represented by 2-19, 2-16, 2-3 and 2-24 did not show neutralization at up to 37.5 µg/ml.

The psNbs were further classified by epitope binning using size exclusion chromatography (SEC). psNb-RBD_SARS-CoV-2_ complexes were competed with high-affinity benchmark Nbs (Nb21, Nb105, and Nb36) that bind distinct and well-characterized epitopes (*16*) **(fig. S3, A and B, Methods**). psNbs fall into five groups: group A (3%) competes with Nb21 and targets RBS, group B (33%) with Nb105, group C (9%) with Nb36, and group D (17%) does not compete with any benchmark Nbs. Approximately one third of psNbs (Group E, 38%) bind strongly to RBDs based on ELISA, but do not neutralize pseudotyped SARS-CoV-2 efficiently. We selected a high-affinity psNb (2-47) from group E and confirmed its strong binding to the fully glycosylated but not the de-glycosylated RBD (**fig. S3C, Methods**). It is possible that 2-47 can target an epitope encompassing the conserved, glycosylated RBD residue(s) (e.g., N343) (*17*). Together, these experiments suggest that a large cohort of high-affinity and divergent psNbs targeting multiple dominant RBD epitopes contribute to the broad-spectrum serologic activities.

17 psNbs spanning four SEC groups (A-D) were tested for binding to RBDs of five SARS-CoV-2 variants, including the fast-spreading Omicron, and other 18 different sarbecovirus RBDs. These psNbs bind strongly to all variants that were evaluated. 16 of the 17 psNbs bind to all four clades (**Fig. 2A**). Interestingly, seven psNbs (such as 2-31 and 2-45) have exceptionally broad activities and bind strongly to all 24 RBDs with a median ELISA IC_50_ of 3 nM (**Fig. 2A, fig. S4, table S1**). Two representative psNbs were measured for binding kinetics to all four clades by SPR. The K_D_s of 2-31 for RBD_SARS-CoV-2_ (Clade 1b), RBD_SARS-CoV_ (clade 1a), RBD_RmYN02-CoV_ (clade 2), RBD_BM-4831-CoV_ (clade 3) are <1 pM, 3.96 pM, 0.59 nM and 3.60 nM, respectively(**Fig. 2D, fig. S7, A and B**). The binding of 2-45 is equally strong for clades 1b, 2 and 3 (0.39 nM, 1.4 nM and 44 pM, respectively) and more moderate for clade 1a (304 nM) (**Fig. 2E, fig. S7, C and D**). Of note, these psNbs are highly specific to sarbecoviruses and do not cross-react with a human whole protein extract at high concentrations (up to 8 µM) (**Fig. 2A**).

**Fig. 2.**
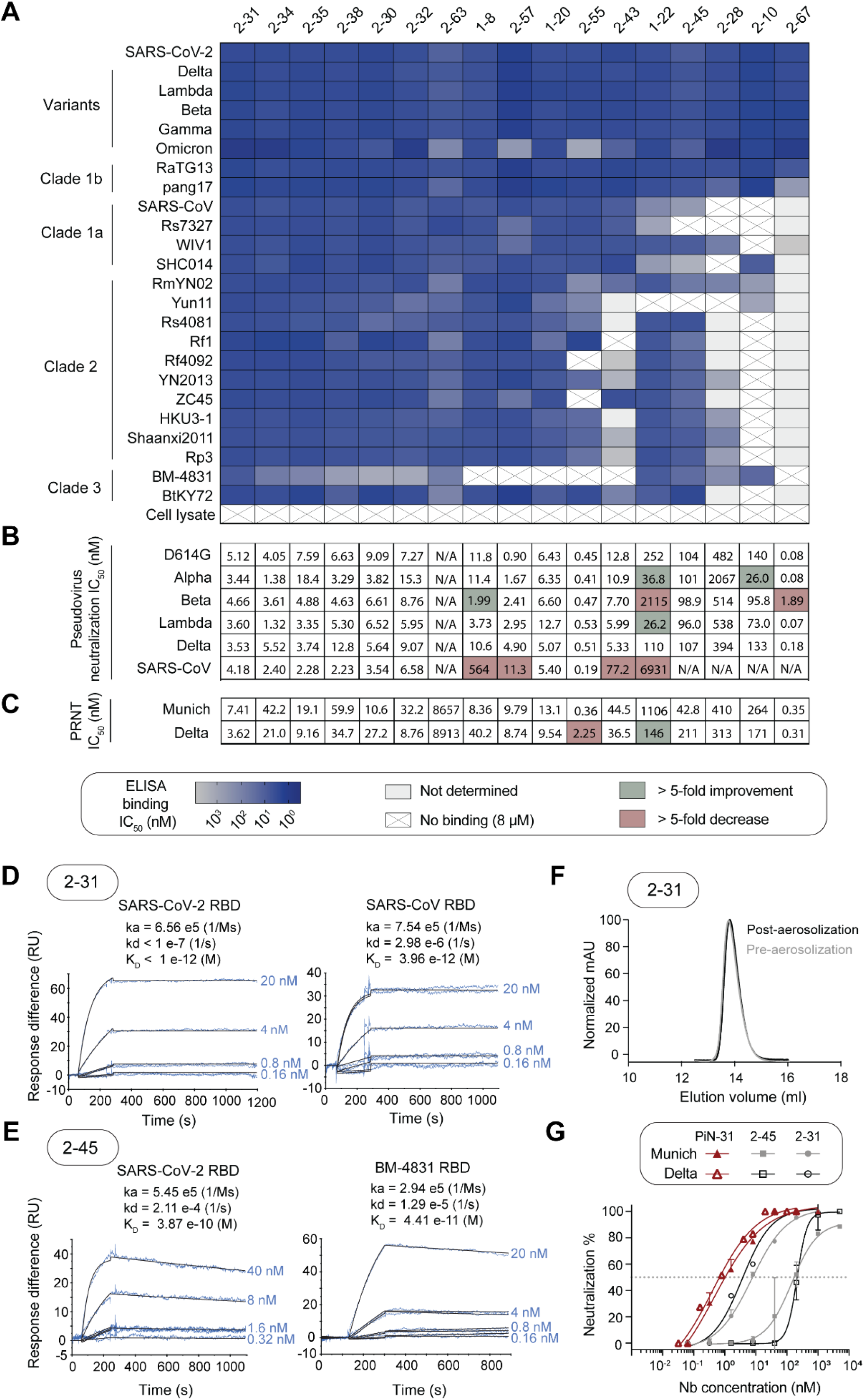
Binding and neutralization of 17 psNbs and development of a bispecific and inhalable psNb (PiN-31). **(A)** Heatmap summary of psNbs for binding to different RBDs by ELISA. White with cross mark: no binding. Silver: binding signals were detected at 8 μM yet the IC_50_s have not been determined. Blue shades: IC_50_ between 0.8 nM to 8 μM. **(B)** Neutralization potency of psNbs against pseudotyped SARS-CoV-2, its variants and SARS-CoV. N/A: no neutralization detected at the highest Nb concentration (2.5 µM). **(C)** Neutralization potency against SARS-CoV-2 Munich strain and the Delta VOC by the PRNT assay. **(D)** Binding kinetic measurements of 2-31 for RBDSARS-CoV-2 and RBDSARS-CoV by surface plasmon resonance (SPR). **(E)** SPR measurements of 2-45 for RBDSARS-CoV-2 and RBDBM-4831-CoV. **(F)** SEC analysis of 2-31 before and after aerosolization with soft mist inhaler. **(G)** Neutralization potency of a bispecific psNb construct (PiN-31) by the PRNT assay.

All but one of these 17 psNbs can potently inhibit SARS-CoV-2 and VOCs *in vitro*, based on a pseudovirus assay, and plaque reduction neutralization test (PRNT) against the Munich strain and Delta VOC (**Fig. 2B and C**, **figs. S5 and S6, Methods**). Group A Nb 2-67, which preferentially binds to clade 1b, is extremely potent for SARS-CoV-2 and VOCs with a median neutralization potency of 1.2 ng/ml. Group C (1-22, 2-10 and 2-45) neutralizes VOCs at single-digit µg/ml, but is less effective against SARS-CoV. 2-63 (group D) has exceptional breath while the other group D psNb 2-28 has limited breadth. Both weakly neutralize SARS-CoV-2 (IC_50_ 7- 132 µg/ml). The remaining psNbs belong to group B and show comparably high potency against SARS-CoV. The median potency for SARS-CoV-2 and VOCs is 86 ng/ml with the most potent (2-55) at 6 ng/ml. Notably, psNbs are usually highly stable and can withstand aerosolization without compromising activity as exemplified by 2-31 and 2-45 (**Fig. 2F, fig. S7, E and F1**, **Methods)**. To demonstrate the high bioengineering potentials of psNbs, we developed a bispecific construct (PiN-31) by fusing these two potent and broad-spectrum psNbs, which cover two distinct epitopes based on SEC. Compared to the monomers, the potency of PiN-31 improves by an order of magnitude to 0.4 nM based on the PRNT assay (**Fig. 2G**).

### Diversity, convergence and evolution of broadly neutralizing psNbs

To understand the mechanisms of broad neutralization, we implemented cryoEM to interrogate the structures of 11 psNbs in complex with the SARS-CoV-2 spike or RBD (**table S2**). Additionally, two atomic psNb: RBD structures were determined by X-ray crystallography (**table S3**). Epitope clustering based on high-resolution structures revealed five primary and distinct epitope classes in RBD_SARS-CoV-2_ (**Fig. 3A**). None of our psNbs (except for 2-57) overlap with key RBD mutations in variants including Alpha, Beta, Delta, Lambda, Gamma, and Omicron (**Fig. 3B**), in contrast to most other Nbs that have been structurally characterized (*18–25*)(**fig. S8**). Over three-quarters of the solvent-exposed and highly conserved RBD residues (sequence identity > 85%) are covered by psNbs (see **Methods**). Superposition onto the spike structure reveals that psNbs preferentially lock RBDs in the 3-up conformation (**Fig. 3C**). Despite having different epitopes and orientations, the small size of Nbs facilitates simultaneous binding of three copies to the spike trimer in distinct and highly symmetric configurations.

**Fig. 3.**
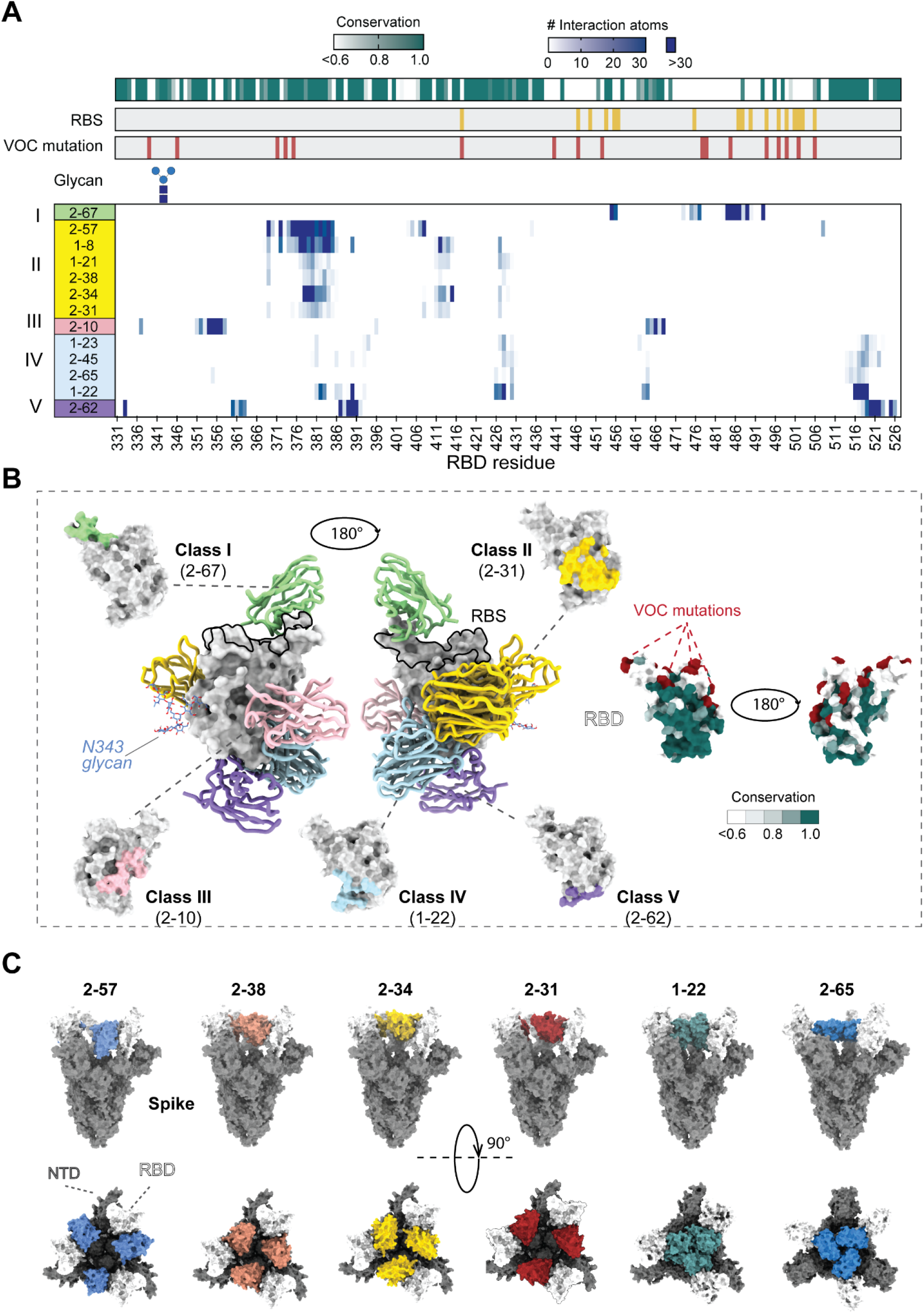
Summary of broadly neutralizing RBD epitopes and spike conformations. **(A)** Clustering analysis of psNbs epitopes. RBS residues are in yellow. The RBD glycosylation site (N343) is denoted. Sarbecovirus sequence conservation is shown in green gradient. The number of psNb: RBD interaction atoms per epitope residue is shown in blue shades. **(B)** Structural representations of five classes of psNbs in complex with RBD and the corresponding epitopes. Light green: class I (2-67), gold: class II (2-31), pink: class III (2-10), light blue: class IV (1-22) and purple: class V (2-62). VOC residues are in red. RBD glycosylation (N343) is in cornflower blue. The same set of perspective view was used for comparison between psNb epitopes and VOC mutation sites. **(C)** Structural diversity of psNbs in complex with the SARS-CoV-2 pre-fusion spike glycoprotein (by cryoEM). RBDs are shown in white, and non-RBD regions of spike in grey.

The **Class II** psNbs are overrepresented in our collection. Phylogenetic analysis reveals that while psNbs are diverse, those isolated from the booster converge more than the initial bleed (**Fig. 4A**). Two subclasses within the class II (A and B) differ in epitopes and Nb binding orientation (**Fig. 4, B and D**). Nevertheless, II(A) and II(B) psNbs share a conserved hydrophobic core epitope (aa 377-386) (**Fig. 4C**) containing two bulky hydrophobic residues (F377 and Y380), and this region is stabilized by a disulfide bond (C379-C432). The improved breadth in class II(B) psNbs, especially for clade 3 RBD_BM-4831-CoV_ lies in its ability to target an additional conserved region with peripheral charged residues (aa 412-415, 427-429). In contrast, subclass II(A) psNbs bind to a less conserved epitope that is composed of primarily hydrophobic residues (e.g., L368, A372, and Y508) (**Fig. 4C**). Notably, A372 on SARS-CoV-2 (clade 1a) is substituted to S/T in other clades, forming a consensus glycosylation motif (^370^NST/S^372^) (*26*), which may explain the reduced neutralization potency of class II (A) psNbs versus class (B) psNbs against SARS-CoV (**Fig. 2B, 4B**).

**Fig. 4.**
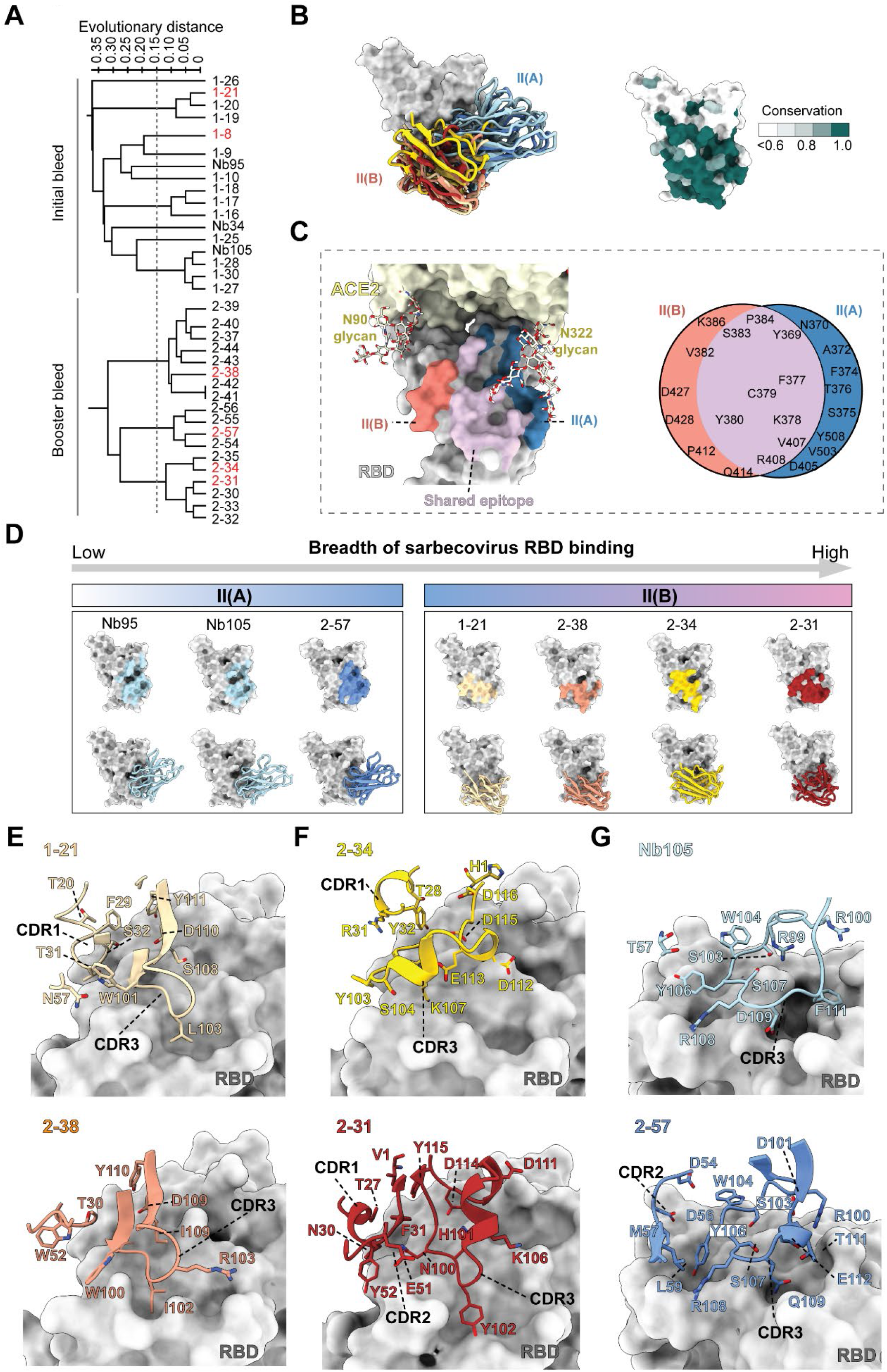
Structural diversity, convergence and evolution of class II psNbs. **(A)** Phylogenetic analysis of psNbs from SEC group B. psNbs with structures determined in this study are in red. Scale of the evolutionary distance: 0-0.35. **(B)** Superposition of class II psNbs reveals two subclasses II(A) and II(B). **(C)** Overlap between II(A) and II(B) epitopes. Purple: shared epitope; blue: non-overlapping epitope of II(A); salmon: non-overlapping epitope of II(B); light yellow: hACE2 with modeled glycans (N90 and N322). **(D)** Structural overview of class II psNbs in complex with RBD and their epitopes. Light blue: Nb95 and Nb105; cornflower blue: 2-57; beige: 1-21; salmon: 2-38; gold: 2-34; dark red: 2-31. **(E-G)** Structural comparison of different Nb pairs (e) 1-21 and 2-38, (f) 2-34 and 2-31, and (g) Nb105 and 2-57 for RBD binding.

**Class II(B)** psNbs comprise two large clusters possessing the best breadth and potency (**Fig. 1D**). While the epitopes largely overlap in the complex structure, the molecular interactions vary substantially. For example, two related psNbs (2-38 and 1-21) are characterized by a short CDR3 (13 aa) forming a β-strand hairpin conformation. Yet, psNb 2-38 is an all-clade binder whereas 1-21 binds strongly to all but RBD_BM-4831-CoV_ (clade 3). The difference in binding is likely due to the K378Q substitution in the RBD, which disrupts a critical salt bridge between D110 (1-21) and K378 (RBD_SARS-CoV-2_) (**Fig. 4E, fig. S9B, S12, A and B**). psNbs 2-34 and 2-31 (from the other dominant cluster) bind all 24 RBDs strongly with potent antiviral activities. The interactions are predominantly mediated by a long CDR3 loop (18 aa) with a small helix. These Nbs form hydrophobic interactions and also bind strongly to conserved charged residues (such as D427 and R408) through electrostatic interactions (**Fig. 4F, fig. S9A, S12, C and D**).

**Class II(A)** psNb (2-57) shares high sequence similarity with Nb105 (isolated from the initial bleed). The major difference lies in their CDR3 heads. Here, a small CDR3 loop of ^109^DLF^111^ in Nb105 is substituted by ^109^QST^111^ in 2-57, which inserts into two conserved pockets (RBD residues F377,Y369,F374 and residues S375,T376,Y508,V407) forming extensive hydrophobic interactions with the RBD (**Fig. 4G, fig. S10, S11, S12E**). This substitution allows psNb 2-57 to tolerate sequence variation better, exhibiting over 100 fold higher affinities for clade 2 RBDs (**table S1**).

**Class I** psNbs that bind the RBS are extremely potent yet rare (∼3%). The structure of the ultrapotent 2-67 was resolved by cryoEM in complex with the RBD (**fig. S13**). Superposition of 2-67:RBD and hACE2:RBD (PDB 6M0J) (*27*) structures revealed major clashes between all three Nb framework regions and two CDRs (2 and 3) with hACE2 (aa 26-37, 66-87, 91-109, 187-194) (**Fig. 5A**). 2-67 targets the protruding RBD ridge (aa 472-490) (**Fig. 5B, fig. S15A**). Binding energy analysis (**Methods**) reveals that compared to other RBS Nbs (*18–25*), the RBD binding epitope of 2-67 is minimally contributed by critical VOC residues (**Fig. 5C, table S5**). Here, 2-67 utilizes all CDRs to form extensive networks of hydrophobic and nonpolar interactions with two critical RBD residues that are not present as VOC mutations (F486 and Y489), and therefore, can achieve ultrahigh affinity for SARS-CoV-2 and retain strong binding and neutralization against the variants (**Fig. 5D**). Moreover, 2-67 can outcompete ultrahigh-affinity Nb21 for RBS binding, thereby further corroborating the ultra-high affinity of this interaction (**fig. S14A**) (*14*). Interestingly, quantitative mass spectrometric analysis reveals that ultrapotent RBS binders (e.g., Nbs 20 and 21) identified from the initial bleed were absent in the booster, implying that they are sourced from short-lived plasma cells (*28*) (**fig. S14B**).

**Fig. 5.**
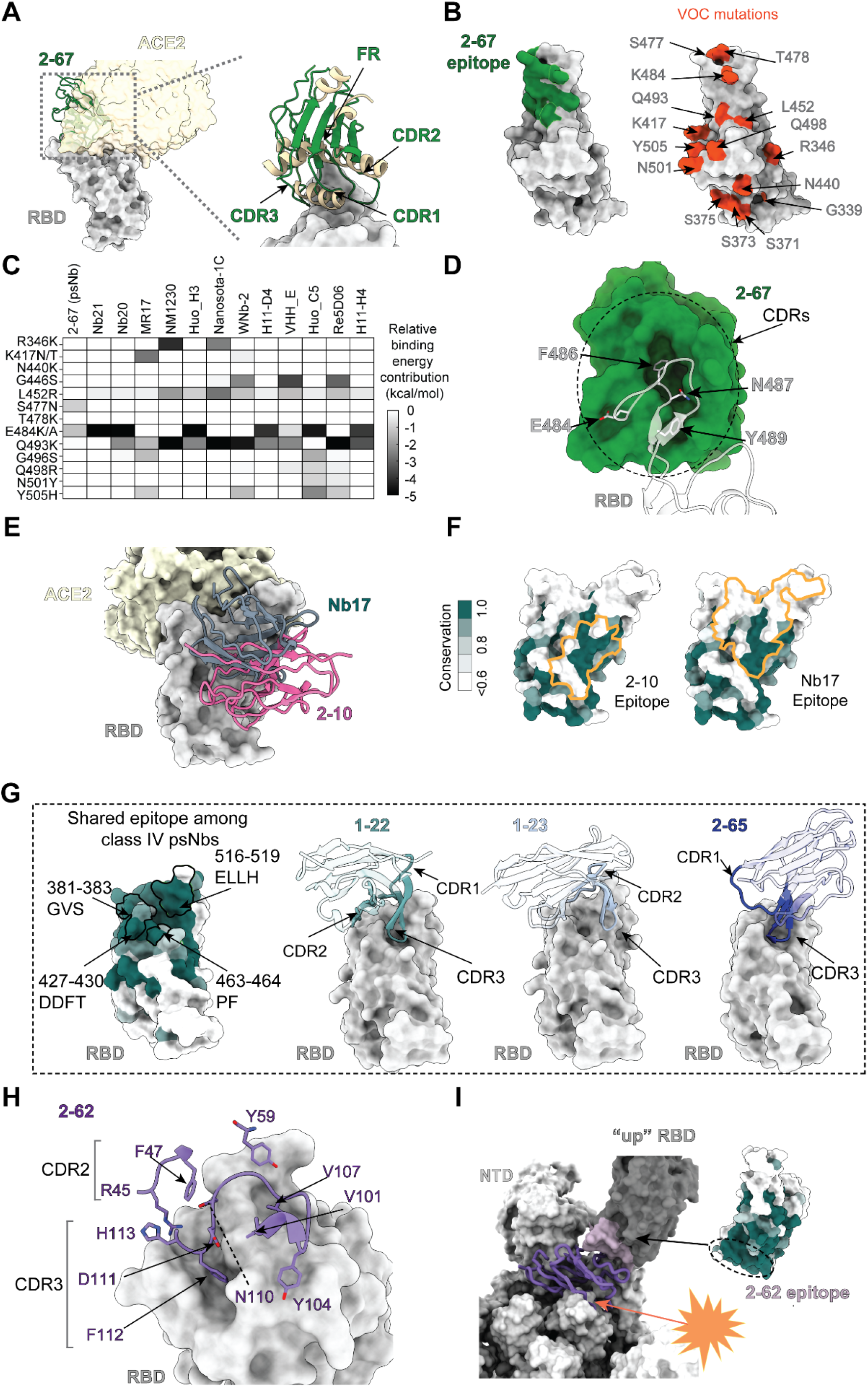
Structural insights of four classes of psNbs. **(A)** Superposition of a class I psNb (2-67): RBD with hACE2 showing steric effects. **(B)** 2-67 epitope (green) and critical VOC mutations (red) on RBD. **(C)** Heatmap of relative binding energy contribution of Nbs per epitope residue that overlaps with critical mutations from VOCs. **(D)** Surface representation showing all CDRs of 2-67 with RBD. Green: 2-67; white: RBD. **(E)** Superposition of a Class III psNb (2-10) and a non-psNb (Nb17). **(F)** The conservation map of RBD and the epitopes of a Class III psNb (2-10) and Nb17. **(G)** Shared epitopes and binding of three class IV psNbs (1-22, 1-23 and 2-65) to the RBD. Shared epitope residues are highlighted on the RBD structure with sequences labeled on the side. **(H)** Binding of a class V (2-62) with the RBD. Interacting CDR residues are labeled. **(I)** Superposition of 2-62 from the 2-62:RBD structure showing the clash between 2-62 and the spike in the all-RBD-up conformation. NTD: N-terminal domain. Representation showing the 2-62 epitope is conserved but not accessible on the spike.

**Class III** psNb (2-10) can destabilize the spike *in vitro* as we observed previously (*16*). However, the structure can be resolved by reconstituting an RBD: 4 Nbs complex by cryoEM (**Methods, fig. S13**). 2-10 targets a rare non-RBS epitope that partially overlaps with that of Nb17 (**Fig. 5E**) isolated from the initial bleed that is partially effective against variants (*16*). Class III psNb binds strongly to RBD variants and clade 1b and shows selectivity within other clades. Compared to Nb17 with limited breadth, the improved breadth of class III psNb is contributed by a recognition motif that shifts (from aa 356-357 to aa 449-450,481-484 and 490-494) towards a smaller and more conserved epitope (**Fig. 5F, fig. S15B**).

Four **Class IV** psNbs (1-22, 1-23, 2-65 and 2-45) were resolved by cryoEM. Their binding involves a plethora of molecular contacts that are dependent upon distinct scaffold orientations, different CDR combinations and sequence-specific bonding networks. Class IV shares a highly conserved and cryptic epitope that is only accessible in the RBD-up conformation (**Fig. 3, A and C, Fig. 5G**). The epitope is characterized by charged and hydrophobic residues (D427/D428, T430, F464, _516_ELLH_519_) forming a cavity (**Fig. 5G**). CDR3s of class IV adopt a “hairpin” conformation that inserts into the RBD cavity forming extensive hydrophobic and polar interactions. 1-22 and 1-23 are all-clade binders that recognize multiple other conserved RBD residues (L390, P426 and P463 for 1-22 and Y369 for 1-23) through hydrophobic interactions (**fig. S15, C and D**). 2-65 shows selectivity towards clade 1b and clade 3 likely due to a unique salt bridge between D124 (CDR3) and the semi-conserved H519 on RBDs (**fig. S15E**). Notably, the class IV epitope partially overlaps with a newly discovered broadly neutralizing antibody S2H97(*10*) and targets RBD from a distinct angle (**fig. S16**). Compared to S2H97 which depends on bulky aromatic side chains (W, F, and Y) for interaction, the binding of 1-22 is primarily mediated through a combination of hydrophobic, non-aromatic (L, I, and V), and basic (R and H) residues. Facilitated by a unique orientation, **Class V** psNb (2-62, which shares high CDR similarity with 2-63) targets RBD through a conserved epitope (T333,L387,_389_DLC_391_, C525, L517, and P521) completely buried in the spike and a region that marginally overlaps with the binding motif of class IV psNbs (i.e., L517 and H519) (**Fig. 5, H and I**). While 2-62 binds tightly to the RBD, it does not bind the pre-fusion spike and is therefore a poor neutralizer of the virus (**fig. S6**).

### Mechanisms of broad neutralization by psNbs

Our comprehensive investigation provides direct evidence that with successive immunization, antibodies emerged in the serum that concurrently and almost exhaustively target diverse and conserved physicochemical and geometric sites on the RBD. We observed that ultrahigh-affinity Nbs (i.e., sub-nM) exhibit neutralization potencies that correlate inversely, almost perfectly, with distance of their epitopes from the RBS (**Fig. 6, A and B**, **Methods**). Ultrapotent class I psNbs neutralize the virus at single-digit ng/ml by binding to the RBS to block hACE2 binding (**fig. S17**). Class II psNbs can still efficiently neutralize the virus (i.e., single-digit ng/ml) primarily by sterically interfering with the receptor binding. In particular, binding to the RBD can clash with the glycan moiety at N322 of hACE2, especially bulky complex-type glycans (*29*) (**fig. S17**). Class III, IV and V possess substantially weaker potencies. Their epitopes are distant from RBS and do not compete with hACE2 *in vitro*.

**Fig. 6.**
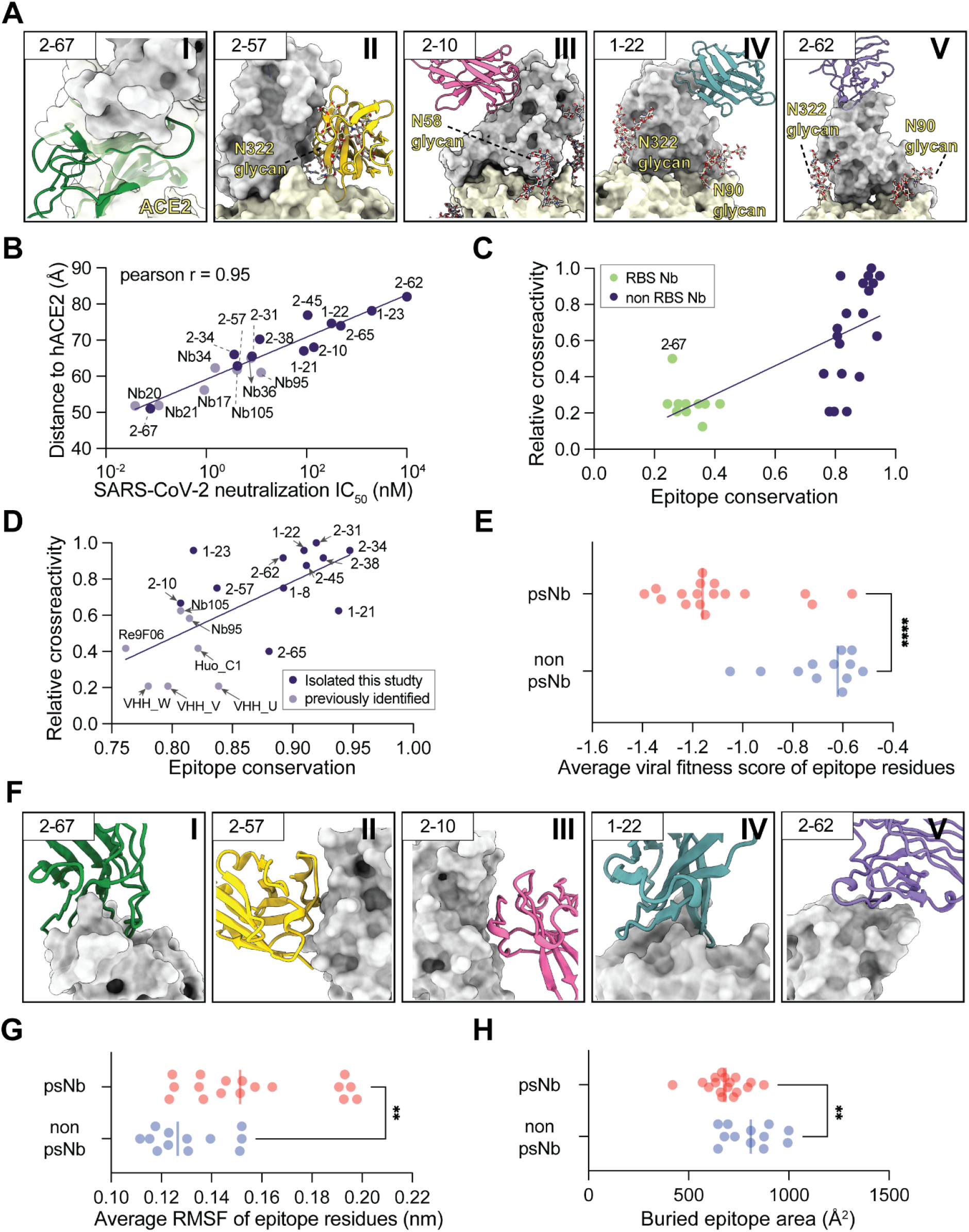
Mechanisms of broad neutralization. **(A)** Superpositions of different psNb classes with RBD and hACE2. Key hACE2 glycosylation sites are shown (N90 and N322). **(B)** Correlation between neutralization potency and the distance between Nb epitope and RBS. The distance is calculated based on centroids of the Nb epitope and hACE2 epitope. **(C–D)**.Correlation between cross-reactivity and epitope conservation for c) all RBD Nbs and d) non-RBS Nbs. The relative cross-reactivity is represented by the normalized ELISA binding of 4 sarbecovirus RBD clades (**table S4**). **(E)** Comparison of the averaged viral fitness score of RBD epitope residues between psNbs and non-psNbs. The fitness score (*31*) is obtained by evaluating the mutational effects on RBD expression level. Negative values correspond to higher loss of fitness. **(F)** Structures showing the geometric features of Nb:RBD interfaces for 5 classes of psNbs. **(G)** Comparison of average root-mean-square-fluctuation (RMSF) of epitope residues between psNbs and non-psNbs (**table S5**). **(H)** Comparison of buried surface area of epitope between psNbs and non psNbs.

To better understand the broad activities of psNbs, we also expressed a set of high-affinity RBD Nbs that have been previously characterized (*18–25*) and evaluated cross-reactivity by ELISA (**table S4, Methods**). Nbs fall into two groups based on RBS occupancy. Overall, their cross-reactivities positively correlate with epitope sequence conservation (**Fig. 6C**). psNbs that do not bind the RBS can be separated based on high epitope conservation (>0.85) (**Fig. 6D**). Critical mutations from the variants are predominantly located on the RBS but also recently on a small patch of relatively conserved residues (e.g., aa 371, 373 and 375) in Omicron (B.1.1.529). Additionally, the virus can mutate on A372S/T to acquire an N-linked glycan (N370) similar to other sarbecoviruses to blunt the host antibody response (*30*). While most Nbs and human IgGs directly interact with these residues, psNbs are barely affected (**Fig. 2A-B**, **fig. S5**). Mutations on these highly conserved epitopes can cause RBD instability reducing viral fitness (*31*) (**Fig. 6E**).

Compared to RBS Nbs that bind to concave epitopes, psNbs generally target smaller (**Fig. 6H**) and flat (class II,III and V) or convex (class I and IV) regions (**Fig. 6F**). psNb epitopes are also more flexible based on average root-mean-square-fluctuation (RMSF) for epitope residues (**Fig. 6G, table S5, Methods**). Together, these properties may render high-affinity binding particularly challenging. Consistently, ScanNet, a geometric deep-learning model reveals that RBS epitopes are predominantly targeted by Nbs (and IgG antibodies) while the psNb epitopes are hardly recognized (**fig. S19**). Moreover, compared to non-psNbs, psNbs utilize almost exclusively hypervariable CDR loops (**fig. S18**). Conceivably, extensive affinity maturation is required to bind these conserved yet difficult-to-bind epitopes.

## Discussion

SARS-CoV-2 continues to evolve, producing variants with high transmissibility and potential to evade host immunity (*32, 33*). New evidence indicates that infection or vaccination by boosting can improve the host antibody response against SARS-CoV-2 variants (*3–7*). Here we discovered that successive immunization of a camelid by recombinant RBD can enhance the development of super-immunity. Integrative proteomics facilitated rapid identification of a large repertoire of high-affinity V_H_H antibodies (Nbs) from the immunized sera against SARS-CoV-2 VOCs and the full spectrum of sarbecovirus family. CryoEM and X-ray crystallography were able to systematically map broadly neutralizing epitopes and interactions, providing insights into the structural basis and evolution of serologic activities. Our data support the notion that RBD structure alone can drive this impressive level of evolution, reshaping the immunogenic landscape towards conserved epitopes. The initial immune response predominantly targets the RBS due to its favorable properties for protein-protein interactions (both host receptor and antibody binding). High-affinity antibodies can evolve to saturate this critical region with extremely high neutralization potency. However, broadly neutralizing antibodies will emerge (with unprecedented diversity) and can continuously develop improved affinity to target conserved epitopes, which provide resistance to antigenic drift. Their neutralization potencies vary substantially yet are strongly and negatively correlated with the distance of epitopes to the RBS-center for viral entry and effective host immunity. Together, our findings inform on the development of safe and broadly protective countermeasures such as vaccines and therapeutics.

Preference for non-conserved epitopes is also observed in other viral antigen structures (**fig. S20**). We suggest that evolutionary conservation imposes constraints on spatio-chemical surface properties, which in turn constraints the immunogenicity of and access to epitopes, as hinted by an overall negative correlation between conservation and predicted epitope propensity (**fig. S21**). However, antibody repertoires are dynamic and can evolve towards conserved epitopes. In addition to COVID-19 super-immunity, potent broadly neutralizing antibodies have also been isolated from HIV elite controllers, although they usually take years to evolve (*34*).

Our highly selected psNbs can bind strongly and specifically to all sarbecoviruses for potent neutralization with the best median antiviral potencies at single-digit ng/ml, which is extremely rare for cross-reactive antibodies (*35*). Multivalent constructs (such as PiN-31) that target distinct and conserved epitopes and their cocktails may provide comprehensive coverage against future SARS-CoV-2 antigenic drift and new sarbecovirus challenges. The low production costs and marked stability of Nbs (*19, 23, 36–38*) (and other miniproteins (*39*) allow for more equitable and efficient distributions globally, particularly for developing countries and regions that are vulnerable to viral spillovers. Combined with their small size (high effective dose and bioavailability) and flexible administration routes that protect both upper and lower respiratory tracts to limit airborne transmissions (*40*), ultrapotent and inhalable psNbs are therefore highly complementary to vaccines, small molecule drugs, and monoclonal antibody therapeutics (*41, 42*). The prospects of winning the race against future outbreaks will rely on the fast development and equitable distribution of an arsenal of broadly protective, cost-effective and convenient technologies.

## ACKNOWLEDGEMENT

We thank P.J. Bjorkman (Caltech) for sharing plasmids bearing sarbecovirus RBDs, the UPMC Genome Center for Illumina MiSeq, and Yong Joon Kim for proofreading of the manuscript. We thank Wenli Yu, Henry Tien, Xueyong Zhu, Meng Yuan, and Robyn L. Stanfield, from the Wilson lab for help with insect cell culture, crystal screening, and X-ray data collection.

## Author contributions

Y.S. conceived the study. Y.X. identified and characterized Nbs. Z.S. developed scripts and analyzed structures. W.H. and D.J.T. solved cryoEM structures. H.L. and I.A.W. determined and analyzed X-ray crystallographic structures. J.T. and D.S. analyzed viral antigenicity. S.N. and P.W.D performed the PRNT assay. Y.S and Y.X. drafted the manuscript with substantial input from I.A.W., W.H., Z.S., D.S., H.L., and D.J. T. All authors reviewed the manuscript. We thank D.S.Reed (University of Pittsburgh) and N.A.Crossland (Boston University) for technical assistance.

## Data and material availability

Plasmids of psNbs can be requested from Y.S.

## Supplementary Materials for

### Materials and Methods

#### Purification of recombinant sarbecovirus RBDs and SARS-CoV-2 spike

The mammalian expression vectors (*43*) encoding the RBDs of RaTG13-CoV (GenBank QHR63300; S protein residues 319-541), SHC014-CoV (GenBank KC881005; residues 307-524), Rs4081-CoV (GenBank KY417143; residues 310-515), pangolin17-CoV (GenBank QIA48632; residues 317-539), RmYN02-CoV (GSAID EPI_ISL_412977; residues 298-503), Rf1-CoV (GenBank DQ412042; residues 310-515), W1V1-CoV (GenBank KF367457; residues 307-528), Yun11-CoV (GenBank JX993988; residues 310-515), BM-4831-CoV (GenBank NC014470; residues 310-530), BtkY72-CoV (GenBank KY352407; residues 309-530) with an N-terminal Mu phosphatase signal peptide and C-terminal His-tag were a kind gift from Pamela J. Bjorkman’s lab, Caltech. Plasmids of the RBDs for the following sarbecovirus strains were synthesized from Synbio Technologies in a similar way: SARS-CoV-2 (GenBank MN985325.1; S protein residues 319-539), SARS-CoV (GenBank AAP13441.1; residues 318-510), Rs7327-CoV (GenBank KY417151.1; residues 319-518), Rs4092- CoV (GenBank KY417145.1; residues 314-496), YN2013-CoV (GenBank KJ473816.1; residues 314-496), ZC45-CoV (GenBank MG772933.1; residues 327-509), HKU3-1-CoV (GenBank DQ022305.2; residues 322-505), Shaanxi2011-CoV (GenBank JX993987.1; residues 321-502) and Rp3-CoV (GenBank DQ071615.1; residues 322-504). The cDNA encoding SARS-CoV-2 spike HexaPro (S) was obtained from Addgene (Cat# 154754) (*44*). To express the proteins, Expi293F cells were transiently transfected with the plasmid using the ExpiFectamine 293 kit (Thermo, Cat#A14635). After 24 hrs of transfection, enhancers were added to further boost protein expression. Cell culture was harvested 5-6 days after transfection and the supernatant was collected by high-speed centrifugation at 21,000×g for 30 min. The secreted proteins in the supernatant were purified using His-Cobalt resin (Thermo). Eluted proteins were then concentrated and further purified by size-exclusion chromatography using a Superose 6 10/300 (for S) or Superdex 75 column (for RBDs, Cytiva) in a buffer composed of 20 mM Hepes pH 7.5 and 150 mM NaCl. SARS-CoV-2 RBD variants were obtained from the Acro Biosystems.

#### Successive camelid immunization with RBD and the proteomic identification of psNbs

A llama (“Wally”) was immunized with an RBD-Fc fusion protein (Acro Biosystems, Cat#SPD-c5255) at a primary dose of 0.2 mg (with complete Freund’s adjuvant), followed by three consecutive boosts of 0.1 mg every 2 weeks. The initial bleed was collected 10 days after the final boost as previously described (*14*). Three weeks after the collection of the initial bleed, the llama was immunized again by four consecutive boosts every 2 weeks. The booster bleed was collected 10 days after the final boost. The immunization procedures were performed by Capralogics, Inc. following the IACUC protocol.

To isolate V_H_H antibodies, plasma was first purified from the immunized bleeds by the Ficoll gradient (Sigma). Polyclonal V_H_Hs mixtures were then isolated from the plasma using a two-step purification protocol (*45*). RBDs from 4 sarbecovirus clades (RBDSARS-CoV-2, RBDSARS-CoV, RBDRmYN02 and RBDBM-4831) were coupled to the CNBr-activated sepharose resin for affinity isolation of RBD-specific V_H_Hs. After binding, RBD-specific V_H_Hs were eluted and proteolyzed as previously described (*14, 15*). Efficiently digested peptides were subjected to proteomics analysis by using the nano-LC 1200 that was coupled online with a Q Exactive™ HF-X Hybrid Quadrupole Orbitrap™ mass spectrometer. The MS data obtained from different RBD-specific V_H_H isolations were analyzed by *AugurLlama* to identify high-affinity Nbs for each RBD (*15*). To facilitate the identification of psNbs, the abundance of Nb CDR3 peptides was quantified across different V_H_H isolations. psNbs (represented by CDR3s) are assigned based on two criteria. 1) they must be classified as high-affinity binders for RBD_SARS-CoV-2_. 2) In addition, they must appear in at least one more RBD isolation sample and are not classified as low affinity binders (*15*).

#### Nb DNA synthesis and cloning

The monomeric Nb genes were codon-optimized and synthesized (Synbio). All the Nb DNA sequences were cloned into a pET-21b(+) vector using EcoRI and HindIII restriction sites. The monomeric Nbs 2-31 and 2-45 were also cloned into a pET-22b(+) vector at the BamHI and XhoI sites for periplasmic expression. To produce a heterodimeric Nb 132-118, the DNA fragment of the monomeric Nb 2-31 was first PCR amplified from the pET-21b(+) vector to introduce a linker sequence and two restriction sites of XhoI and HindIII that facilitate cloning:

Primer1: cccAAGCTTggtggtggtggtagtggtggtggtggtagtggtggtggtggtagtCAaGTTCAACTGGTTGAATCTG;

Primer2: ccgCTCGAGTGCGGCCGCcagtttGCTACTAACGGTAACTT.

The PCR fragment was then inserted into the 2-45 pET-21b(+) vector at the same restriction sites to produce the heterodimer 2-45-(GGGGS)_3_-2-31.

#### Purification of Nbs

Nb DNA constructs were transformed into BL21(DE3) cells and plated on Agar with 50 µg/ml ampicillin at 37 °C overnight. Cells were cultured in an LB broth to reach an O.D. of ∼ 0.5- 0.6 before IPTG (0.5 -1 mM) induction at 16/20°C overnight. Cells were then harvested, sonicated, and lysed on ice with a lysis buffer (1xPBS, 150 mM NaCl, 0.2% TX-100 with protease inhibitor). After cell lysis, protein extracts were collected by centrifugation at 21,000 x g for 10 mins and the his-tagged Nbs were purified by the Cobalt resin (Thermo) and natively eluted with a buffer containing 150 mM imidazole buffer. Eluted Nbs were subsequently dialyzed in a dialysis buffer (e.g., 1x DPBS, pH 7.4 or SEC buffer).

For the periplasmic preparation of Nbs (2-31 and 2-45), cell pellets were resuspended in the TES buffer (0.1 M Tris-HCl, pH 8.0; 0.25 mM EDTA, pH 8.0; 0.25 M Sucrose) and incubated on ice for 30 min. The supernatants were collected by centrifugation and subsequently dialyzed to DPBS. The resulting Nbs were then purified by Cobalt resin as described above.

#### ELISA (enzyme-linked immunosorbent assay)

Indirect ELISA was carried out to evaluate the camelid immune responses of the total single-chain only antibody (V_H_H) to an RBD and to quantify the relative affinities of the psNbs. A 96-well ELISA plate (R&D system) was coated with the RBD protein or the HEK-293T cell lysate at an amount of approximately 3-5 ng per well in a coating buffer (15 mM sodium carbonate, 35 mM sodium bicarbonate, pH 9.6) overnight at 4°C, with subsequent blockage with a blocking buffer (DPBS, v/v 0.05% Tween 20, 5% milk) at room temperature for 2 hours. To test the immune response, the total V_H_H was serially 5-fold diluted in the blocking buffer and then incubated with the RBD-coated wells at room temperature for 2 hours. HRP-conjugated secondary antibodies against llama Fc were diluted 1:75,00 in the blocking buffer and incubated with each well for an additional 1 hour at room temperature. For the initial screening of Nb binding against 4 RBDs, scramble Nbs that do not bind the RBDs were used as negative controls. Nbs were serially 10-fold diluted from 1 μM to 1 nM in the blocking buffer. For the Nb affinity measurements against 24 RBDs, Nbs were serially 4-fold diluted. The dilutions were incubated for 2 hours at room temperature. HRP-conjugated secondary antibodies against the T7-tag were diluted at 1:5,000 or 1:75,00 in the blocking buffer and incubated for 1 hour at room temperature. Three washes with 1x PBST (DPBS, v/v 0.05% Tween 20) were carried out to remove nonspecific absorbances between each incubation. After the final wash, the samples were further incubated in the dark with freshly prepared w3,3′,5,5′-Tetramethylbenzidine (TMB) substrate for 10 mins at room temperature to develop the signals. After the STOP solution (R&D system), the plates were read at multiple wavelengths (450 nm and 550 nm) on a plate reader (Multiskan GO, Thermo Fisher). The raw data were processed by Prism 9 (GraphPad) to fit into a 4PL curve and to calculate logIC_50_.

#### Competitive ELISA with recombinant hACE2

A 96-well plate was pre-coated with recombinant spike-6P at 2 µg/ml at 4°C overnight. Nbs were 5-fold diluted (from 0.2/1/5 µM to 12.8/64/320pM) in the assay buffer with a final amount of 50 ng biotinylated hACE2 at each concentration and then incubated with the plate at room temperature for 2 hrs. The plate was washed by the washing buffer to remove the unbound hACE2. 1:6000 diluted Pierce™ High Sensitivity NeutrAvidin™-HRP (Thermo fisher cat# 31030) were incubated with the plate for 1 hr at room temperature. TMB solution was added to react with the HRP conjugates for 10 mins. The reaction was then stopped by the Stop Solution. The signal corresponding to the amount of the bound hACE2 was measured by a plate reader at 450 nm and 550 nm. The wells without Nbs were used as control to calculate the percentage of hACE2 signal. The resulting data were analyzed by Prism 9 (GraphPad) and plotted.

#### psNb epitope analysis by competitive size exclusion chromatography (SEC)

Analytical size exclusion chromatography was performed with a Superdex 75 increase GL column (column volume: 24 mL, Cytiva) on a Shimadzu HPLC system equipped with a multi-wavelength UV detector at a flow rate of 0.3 mL/min. The column was connected and placed in a column oven set to 15 °C, and the SEC running buffer was 10 mM HEPES pH 7.1, 150 mM NaCl. A reference SEC profile for RBD with epitope I benchmark Nb (Nb21), epitope II benchmark Nb (Nb105) and epitope III benchmark Nb (Nb36) was performed after column equilibration. Subsequently, three separate runs with RBD and specified psNb were performed after mixing with (i) benmark I and benchmark II, (ii) benchmark II and III and (iii) benchmark I and III Nbs. A supershift of the peak, at the same or to the left of the RBD and three Nbs peak in the reference profile, in run (i) but not run (ii) and (iii) sorts a psNb into group C. Similarly, psNb with a supershift in run (ii) but not run (i) and (iii) belongs to group A, and a supershift in run (iii) but not run (i) and (ii) belongs to group B. When supershifts were observed in all three runs, the psNb was sorted into a group D. psNbs were sorted into group E if the supershifts were not detected on SEC.

#### Nb affinity measurement by SPR

Surface plasmon resonance (SPR, Biacore 3000 system, GE Healthcare) was used to measure Nb affinities. RBD proteins were immobilized on the activated CM5 sensor-chip in pH 4.0 10 mM sodium acetate buffer. The surface of the sensor chip was blocked by 1 M Tris-HCl (pH 8.5). For each Nb analyte, a series of concentration dilutions was injected in HBS-EP running buffer (GE-Healthcare), at a flow rate of 20 μl/min for 180 s, followed by a dissociation time of 15 mins. Between each injection, the sensor chip surface was regenerated with the low pH buffer containing 10 mM glycine-HCl (pH 1.5 - 2.0). The regeneration was performed with a flow rate of 30-40 μl/min for 30 - 45 s. The measurements were duplicated, and only highly reproducible data were used for analysis. Binding sensorgrams for each Nb were processed and analyzed using BIAevaluation by fitting with the 1:1 Langmuir model or the 1:1 Langmuir model with the drifting baseline.

#### Pseudotyped SARS-CoV-2 neutralization assay

The 293T-hsACE2 stable cell line (Cat# C-HA101, Lot# TA060720C) and pseudotyped SARS-CoV-2 (Wuhan-Hu-1 strain, D614G, Alpha, Beta, Lambda, Delta and Omicron) particles with luciferase reporters were purchased from the Integral Molecular. The neutralization assay was carried out according to the manufacturers’ protocols. In brief, 3- or 5-fold serially diluted Nbs / immunized V_H_H mixture was incubated with the pseudotyped SARS-CoV-2-luciferase for accurate measurements. At least seven concentrations were tested for each Nb and at least two repeats of each Nb were done. Pseudovirus in culture media without Nbs was used as a negative control. 100 µl of the mixtures were then incubated with 100 µl 293T-hsACE2 cells at 2.5×10e^5^ cells/ml in the 96-well plates. The infection took ∼72 hrs at 37 °C with 5% CO_2_. The luciferase signal was measured using the *Renilla*-Glo luciferase assay system (Promega, Cat# E2720) with the luminometer at 1 ms integration time. The obtained relative luminescence signals (RLU) from the negative control wells were normalized and used to calculate the neutralization percentage at each concentration. Data were processed by Prism 9 (GraphPad) to fit into a 4PL curve and to calculate the logIC_50_ (half-maximal inhibitory concentration).

#### SARS-CoV-2 Munich and Delta plaque reduction neutralization test (PRNT)

Nbs were diluted in a 3- or 5-fold series in Opti-MEM (Thermo). Each Nb dilution (110 μl) was mixed with 110 μl of SARS-CoV-2 (Munich strain) containing 100 plaque-forming units (p.f.u.) or 110 μl of SARS-CoV-2 (Delta strain, BEI Cat# NR55611) containing 50 p.f.u. of the virus in Opti-MEM. The Nb–virus mixes (220 μl total) were incubated at 37°C for 1 h, after which they were added dropwise onto confluent Vero E6 cell (ATCC® CRL-1586™, for Munich) or Vero E6-TMPRSS2-T2A-ACE2 cells (BEI cat# NR- 54970, for Delta) monolayers in the six-well plates. After incubation at 37°C, 5% (v/v) CO_2_ for 1 h, 2 ml of 0.1% (w/v) immunodiffusion agarose (MP Biomedicals) for Munich strain and 0.25% (w/v) immunodiffusion agarose for Delta strain in Dulbecco’s modified eagle medium (DMEM) (Thermo) with 10% (v/v) FBS and 1x pen-strep was added to each well. The cells were incubated at 37°C, 5% CO_2_ for 72 hrs. The agarose overlay was removed and the cell monolayer was fixed with 1 ml/well formaldehyde (Fisher) for 20 min at room temperature. The fixative was discarded and 1 ml/well of 1% (w/v) crystal violet in 10% (v/v) methanol was added. Plates were incubated at room temperature for 20 min and rinsed thoroughly with water. Plaques were then enumerated and the 50% plaque reduction neutralization titer (PRNT_50_) was calculated. A validated SARS-CoV-2 antibody-negative human serum control and a validated NIBSC SARS-CoV-2 plasma control were obtained from the National Institute for Biological Standards and Control, UK) and an uninfected cells control were also used to ensure that virus neutralization by antibodies was specific.

#### Biological safety

All work with SARS-CoV-2 was conducted under biosafety level-3 (BSL-3) conditions in the University of Pittsburgh Center for Vaccine Research (CVR) and the Regional Biocontainment Laboratory (RBL). Respiratory protection for all personnel when handling infectious samples or working with animals was provided by powered air-purifying respirators (PAPRs; Versaflo TR-300; 3M, St. Paul, MN). Liquid and surface disinfection was performed using Peroxigard disinfectant (1:16 dilution), while solid wastes, caging, and animal wastes were steam-sterilized in an autoclave.

#### Aerosolization of PiN using a soft-mist inhaler

Nb was eluted and collected in the SEC running buffer (20 mM HEPES, 150 mM NaCl, pH 7.5) and then concentrated to 0.5 ml (1 mg/ml). Nb was aerosolized by using a portable soft mist inhaler Pulmospray® (Resyca). Around 0.1 – 0.15 ml dead volume was observed in the syringe and connector. The aerosols were collected in a 50 ml falcon tube and SEC analysis was performed as described above.

#### Cryo-electron microscopy data collection and image processing

SARS-CoV-2 HexaPro spike at 1.1 mg/mL was incubated with 1.5-fold molar excess of specified Nbs at room temperature for two hours. 3.5 μL of 1 to 3 dilution sample using HBS buffer with 1% glycerol was applied onto a freshly glow discharged UltraAuFoil R1.2/1.3 grid (300 mesh) and plunge frozen using a vitrobot Mark IV (Thermo Fisher) with a blot force of 0 and 2.5 s blot time at 100% humidity at 4 °C. The cryoEM datasets were collected at either CWRU or PNCC.

For CWRU datasets, movie stacks were recorded using an FEI Titan Krios transmission electron microscope G3i operated at 300 keV and equipped with a Gatan K3 direct electron detector and Gatan BioQuantum image filter operated in zero-loss mode with a slit width of 20 eV. Automated data collection was carried out using serialEM (*46*) at a nominal magnification of 81,000x with a physical pixel size of 1.07 Å/pixel (0.535 Å/pixel at super-resolution) for spike:nanobody complexes and 165,000x with a physical pixel size of 0.52 Å/pixel (0.26 Å/pixel at super-resolution) for RBD:nanobodies complexes. Each movie stack was collected with a dose rate of 18 electron/pixel/s in super-resolution mode and fractionated in 40 frames with two-second exposure, resulting in a total dose of ∼31.4 e/Å^2^ for 81,000x and in 32 frames with 1.5-second exposure, resulting in a total dose of ∼103.4 e/Å^2^ for 81,000x. The number of movies for a specified dataset was listed in **table S2**. The defocus range was set to between -0.5 and -2 μm.

For PNCC datasets, movie stacks were recorded using an FEI Titan Krios transmission electron microscope G3i operated at 300 keV and equipped with a Gatan K3 direct electron detector and Gatan BioContinuum image filter operated in zero-loss mode with a slit width of 20 eV. Automated data collection was carried out using serialEM (*46*) at a nominal magnification of 64,000x with a physical pixel size of 1.329 Å/pixel (0.6645 Å/pixel at super-resolution). The dose rate was determined over a sample hole to calculate the exposure time resulting in a total dose of 40 e/Å^2^ and the exposures were fractionated into a total of 40 frames. The number of movies for a specified dataset was listed in **table S2**. The defocus range was set to between -0.5 and -2 μm.

Image processing was performed on-the-fly using CryoSPARC Live version 3.2 (*47–49*). The particles were automatically picked using the blob picker with 240 Å or 100 Å diameter for spike:nanobody or RBD:nanobodies, respectively. Reference-free 2D classification was performed in streaming with 200 classes and limited maximum resolution to 18 Å. Upon the completion of data collection, the selected particles from the good 2D class averages were subjected for another round of 2D classification with 200 classes, and particles from classes with resolution better than 10 Å and ECA less than 2 were selected for subsequent analysis. These 2D classes were submitted for a “rebalance 2D” job type to trim particles from dominant views. The rebalanced particle set was then used for ab-initio reconstruction to generate the initial volume. 3D refinement was first carried out using non-uniform refinement using ab-initio volume as the reference without mask. To resolve the density for nanobodies, an RBD structure (PDB 6M0J) (*27*) was docked into the cryoEM density and the structural model for nanobody was manually placed into the additional density accounted for nanobodies to ensure the correct orientation. The resulting RBD:nanobody model was used to generate the mask for focused 3D classification in CryoSPARC version 3.3.1 with six classes and target resolution of 6 Å, and PCA was chosen as the initialization mode with the parameter “number of components” set for 2. 3D classes with both RBD and nanobody densities well resolved were selected for sequent new local refinement with the mask around RBD and nanobody. The gold-standard Fourier shell correlation (FSC) of 0.143 criterion was used to report the resolution and the FSC curves were corrected for the effects of a soft mask using high-resolution noise substitution (*50*).

#### Model building and refinement

PDB entry, 7CAK (*51*), was used as the initial model for spike excluding RBD and the missing fragment for residues 621 to 640 was modeled de novo in Coot (*52*). RBD from PDB entry, 6M0J (*27*), was used as the initial model and nanobodies structures were generated using ColabFold (*53*). After assembling individual components into a single PDB file, the models were refined into the composite map using phenix.real_space_refine (*54*). The glycans were built using the carbohydrate module in Coot (*55*). Models were manually corrected in Coot (version 9.6.0) between rounds of read-space refinement in Phenix. All statistics for structural models were reported in **table S2**. Fig. panels depicting cryoEM maps or atomic models generated using ChimeraX (*56*). Maps colored by local resolution were generated using RELION 3.1 (*57*).

#### Calculation for binding energy contribution

Molecular dynamic (MD) simulations were used to generate short 1 ns trajectories for relative binding energy calculation. Input files for MD simulations of SARS-CoV-2 RBD and nanobody complexes (**table S5**) were prepared using tleap (*58*). MD simulations were performed using the NAMD (*59*) and the amber ff19sb (*60*), GLYCAM_06j (*61*), ions with the TIP3P water model (*62*). Proteins were solvated in a cubic water box with a 16 Å padding in all directions. Sodium ions and chloride ions were added to achieve a physiological salt condition of 150 mM. The systems were energy minimized for 10,000 steps to remove bad contacts. Then, the systems were equilibrated with all heavy atoms restrained harmonically and the temperature raised 10 K per 10,000 steps starting from 0 to 300 K using temperature reassignment. After reaching the desired temperature, harmonic restraints were gradually reduced using a scale from 1.0 to 0 with a 0.2 decrements for every 50,000 steps. MD simulations were performed under the NPT ensemble (*63, 64*). Langevin dynamics was used for constant temperature control, with the value of Langevin coupling coefficient and the Langevin temperature set to 5 ps and 300 K, respectively. The pressure was maintained at 1 atm using the Langevin piston method with a period of 100 fs and decay times of 50 fs. A time step of 2 fs was used for all the simulations by using the SHAKE algorithm (*65*) to constrain bonds involving hydrogen atoms.

For each snapshot, every 10 ps of a 1 ns trajectory of SARS-CoV-2 RBD and nanobody complexes, the binding energy of MM/PBSA was calculated using Eqs. (1) and (2) (*66, 67*)

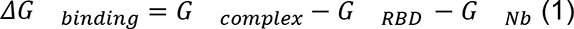

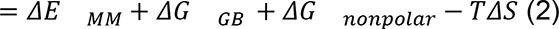

where is the molecular mechanic (MM) interaction energy calculated in gas-phase between RBD and nanobody, including electrostatic and van der Waals energies; the desolvation free energy consists of polar (and nonpolar) terms; is the change of conformational entropy on nanobody binding, which was not considered here as the binding epitope on the RBD is very stable and the comparison was performed internally. The decomposition of the binding free energy to the relative energy contribution from individual residues was performed using the MMPBSA.py module in AMBER18 (*68*). The relative contribution to the binding energy from mutated residues observed in VOC were plotted as heatmap.

#### Crystallographic analysis of psNbs with RBD

The receptor-binding domain (RBD) of the SARS-CoV-2 spike (S) protein (GenBank: QHD43416.1), used in the crystallographic study, was cloned into a customized pFastBac vector (*69*), and fused with an N-terminal gp67 signal peptide and C-terminal His_6_ tag (*70*). Recombinant bacmids encoding each RBDs were generated using the Bac-to-Bac system (Thermo Fisher Scientific) followed by transfection into Sf9 cells using FuGENE HD (Promega) to produce baculoviruses for RBD expression. RBD protein was expressed in High Five cells (Thermo Fisher Scientific) with suspension culture shaking at 110 r.p.m. at 28 °C for 72 hours after the baculovirus transduction at an MOI of 5 to 10. Supernatant containing RBD protein was then concentrated using a 10 kDa MW cutoff Centramate cassette (Pall Corporation) followed by affinity chromatography using Ni-NTA resin (QIAGEN) and size exclusion chromatography using a HiLoad Superdex 200 pg column (Cytiva). The purified protein sample was buffer exchanged into 20 mM Tris-HCl pH 7.4 and 150 mM NaCl and concentrated for binding analysis and crystallographic studies.

1-21+SARS-CoV-2 RBD and 2-31+SARS-CoV-2 RBD+CC12.1 complex were formed by mixing each of the protein components in an equimolar ratio and incubating overnight at 4°C. 384 conditions of the JCSG Core Suite (Qiagen) were used for setting-up trays for screening the 1-21 complex (12 mg/ml) and 2-31 complex (14.3 mg/ml) on our robotic CrystalMation system (Rigaku) at Scripps Research. Crystallization trials were set-up by the vapor diffusion method in sitting drops containing 0.1 μl of protein complex and 0.1 μl of reservoir solution. Crystals appeared on day 3, were harvested on day 12, pre-equilibrated in cryoprotectant containing 0-10% ethylene glycol, and then flash cooled and stored in liquid nitrogen until data collection. X-ray diffraction data were collected at cryogenic temperature (100 K) at beamlines 23-ID-D of the Advanced Photon Source (APS) at Argonne National Laboratory and were collected from crystals grown in drops containing 20% polyethylene glycol 8000, 0.1 M NaCl, 0.1 M CAPS pH 10.5 for the 1-21 complex and drops containing 40% MPD, 0.1M cacodylate pH 6.5, 5% (w/v) polyethylene glycol 8000 for the 2-31 complex. Collected data were processed with HKL2000 (*71*). X-ray structures were solved by molecular replacement (MR) using PHASER (*72*) with original MR models for the RBD and Nanobody from PDB 7JMW (*73*) and PDB 7KN5 (*19*). Iterative model building and refinement were carried out in COOT (*74*) and PHENIX (*75*), respectively.

#### RBD epitope analysis by ScanNet

The epitope propensity profile of SARS-CoV-2 RBD (PDB 7jvb) was computed by ScanNet, a state-of-the-art geometric deep learning model for structure-based protein binding site prediction (*76*). We used the B-cell epitope network (ScanNet-BCE) without evolutionary information that was not trained on any SARS-CoV-1/2 antibody/antigen complex.

#### Conservation and ScanNet analysis of viral antigens

##### Dataset preparation

We collected all viral antibody - antigen complexes in PDB, as listed by SabDab (*77*) (release: 10/19/2021). Antigens were clustered at 70% sequence identity with a minimum length coverage of 15% using CD-HIT (*78*), the Bio.align pairwise sequence alignment module and CATH domain identifiers. Briefly, we found that (i) ngram-based clustering using CD-HIT with default parameters overestimated the number of clusters, while (ii) computing all the entries of the pairwise sequence similarity matrix was intractable for our set of several thousands structures. We instead proceeded as follows: for a given sequence identity cut-off T (e.g. 100), sequences were clustered with CD-HIT, and the resulting representatives were further clustered at T-20. For each pair of T-representatives with identical CATH identifier or same T-20 cluster, the sequence identity and coverage were evaluated by pairwise alignment and the corresponding T-clusters were merged if necessary. The process was iterated at T=100%, 95%, 90%, 70% sequence identity cut-offs and yielded satisfactory clusters. We further grouped together the following antigen clusters that had lower sequence identity but high structure similarity: (i) Influenza hemagglutinin from strains H1N1, H3N2, H5N1, H7N9, H2N2 and (ii) Envelope protein of Dengue 1, Dengue 2, Dengue 4 and Zika. Only clusters with at least 7 unique antibodies were retained for further analysis, yielding 11 viral antigens.

##### Antibody hit rate calculation

We constructed a multiple sequence alignment for each antigen cluster using MAFFT (*79*) and selected a representative structure with highest structure coverage and resolution. For each column of the alignment, the antibody hit rate was calculated as the fraction of unique antibodies binding it. For each complex, we identified the epitope residues as the antigen residues having at least one heavy atom within 4Å of at least one antibody heavy atom. Our analysis was performed on the antigen surface residues (relative accessible surface area >= 0.25, computed within the biological assembly for multimeric antigens using Bio.DSSP). Finally, the column-wise antibody hit rate was projected back onto the surface residues representative structure.

##### Calculation of antigen conservation and epitope propensity

For each representative antigen chain, a multiple sequence alignment was constructed by homology search on the UniRef30_2020_06 sequence database using HHBlits (*80*) (4 iterations, default parameters). The alignment was deduplicated, hits with high gap content (>=25% of the alignment) were discarded and the 10K best hits were retained based on sequence identity. Each sequence was assigned a weight inversely proportional to the number of similar sequences found in the alignment (90% sequence identity cut-off). The amino acid frequency at each site f_i_(a) was subsequently calculated and the residue-wise conservation score was defined as *C_i_ = ln(20) - S(f_i_)* where 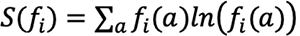. Conservation scores range from 0 to ln(20) = 2.99, higher is more conserved. We checked that this protocol correlated well with the ConSurf method based on phylogenetic trees (*81*). Epitope propensity scores were calculated with ScanNet-BCE.

#### RMSF calculation

Root mean square fluctuation of each RBD residue was calculated based on 100 ns Molecular Dynamics simulation trajectory. The simulation was run starting from the RBD structure (PDB 6lzg) using Gromacs 2020 version with the CHARMM36m force field (*82*). The RBD structure was solvated in transferable intermolecular potential with 3 points (TIP3P) water molecules and ions were added to equalize the total system charge. The steepest descent algorithm was used for initial energy minimization until the system converged at Fmax < 1,000 kJ/(mol · nm). Then water and ions were allowed to equilibrate around the protein in a two-step equilibration process. The first part of equilibration was at a constant number of particles, volume, and temperature (NVT). The second part of equilibration was at a constant number of particles, pressure, and temperature (NPT). For both MD equilibration parts, positional restraints of k = 1,000 kJ/(mol · nm2) were applied to heavy atoms of the protein, and the system was allowed to equilibrate at a reference temperature of 300 K, or reference pressure of 1 bar for 100 ps at a time step of 2 fs. Altogether 10,000 frames were saved for the RMSF analysis at intervals of 10 ps. To estimate average epitope RMSF, we defined epitope residues as residues with at least one atom within 4Å from the Nb atom and averaged RMSF over epitope residues.

#### Phylogenetic analysis

The phylogenetic analysis was performed to 1) 19 Sarbecovirus RBD sequences (**Fig. 1a**) 2) 100 psNb CDR3 sequences (**fig. S2a**) and 3) 32 SEC group B psNb and 3 previously characterized benchmark Nbs (Nb34, Nb95 and Nb105) CDR3 sequences (**Fig. 4a**). Sequences were aligned by MUSCLE (*83*) with default parameters. The phylogenetic tree was then constructed by the MEGA (*84*) using the maximized likelihood estimation method.

#### Sarbecovirus RBD conservation

To calculate Sarbecovirus RBD conservation with respect to SARS-CoV-2, 18 Sarbecovirus RBD sequences mentioned above were aligned to SARS-CoV-2 RBD sequence. After alignment, each SARS-CoV-2 RBD residue was compared to corresponding residue in other RBD sequences. An identical residue to that in SARS-CoV-2 is considered a match. The conservation for each RBD residue is calculated by the number of matches over 18.

#### psNb epitope heat map

An RBD residue and an Nb residue were defined in contact if the distance between any pair of their atoms was lower than a threshold of 4 Å. The Nb contact value of each RBD residue is calculated as the sum of all the Nb contacts.

#### Measurement of buried surface area (BSA)

The solvent-accessible surface area (SASA) of molecules was calculated by FreeSASA(*85*). The buried surface area in the case of the Nb-RBD complex was then calculated by equation:

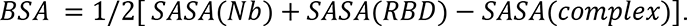

#### Distance between Nb epitope and receptor binding sites

The Nb epitope residues were extracted based on the method described above. The centroid of the Nb epitope residues is defined by the center of mass of atoms. Similarly, the centroid of receptor binding sites(RBS) was extracted. The distance between Nb epitope and RBS was calculated based on the distance between two centroids.

#### Viral fitness score of epitope residues

The viral fitness score was obtained from (*31*) by evaluating the mutational effects on RBD expression level. The fitness score of each RBD residue is the averaged value of mutating to any other amino acids. The viral fitness score of epitopes was calculated by averaging fitness score over epitope residues.

#### Glycosylation modeling

Based on the crystal structure of complex ACE2 and SARS-CoV-2 RBD contain seven (N53, N90, N103, N322, N432, N546, N690) and one (N343) N-glycosylation sites, respectively. A heterogeneous glycosylated system was set up for the complex based on previous study (*29*). The glycosylated ACE2:RBD complex was modeled by the Glycan Modeler module of the CHARMM-GUI webserver (*86*).

---

**Fig. S1.**
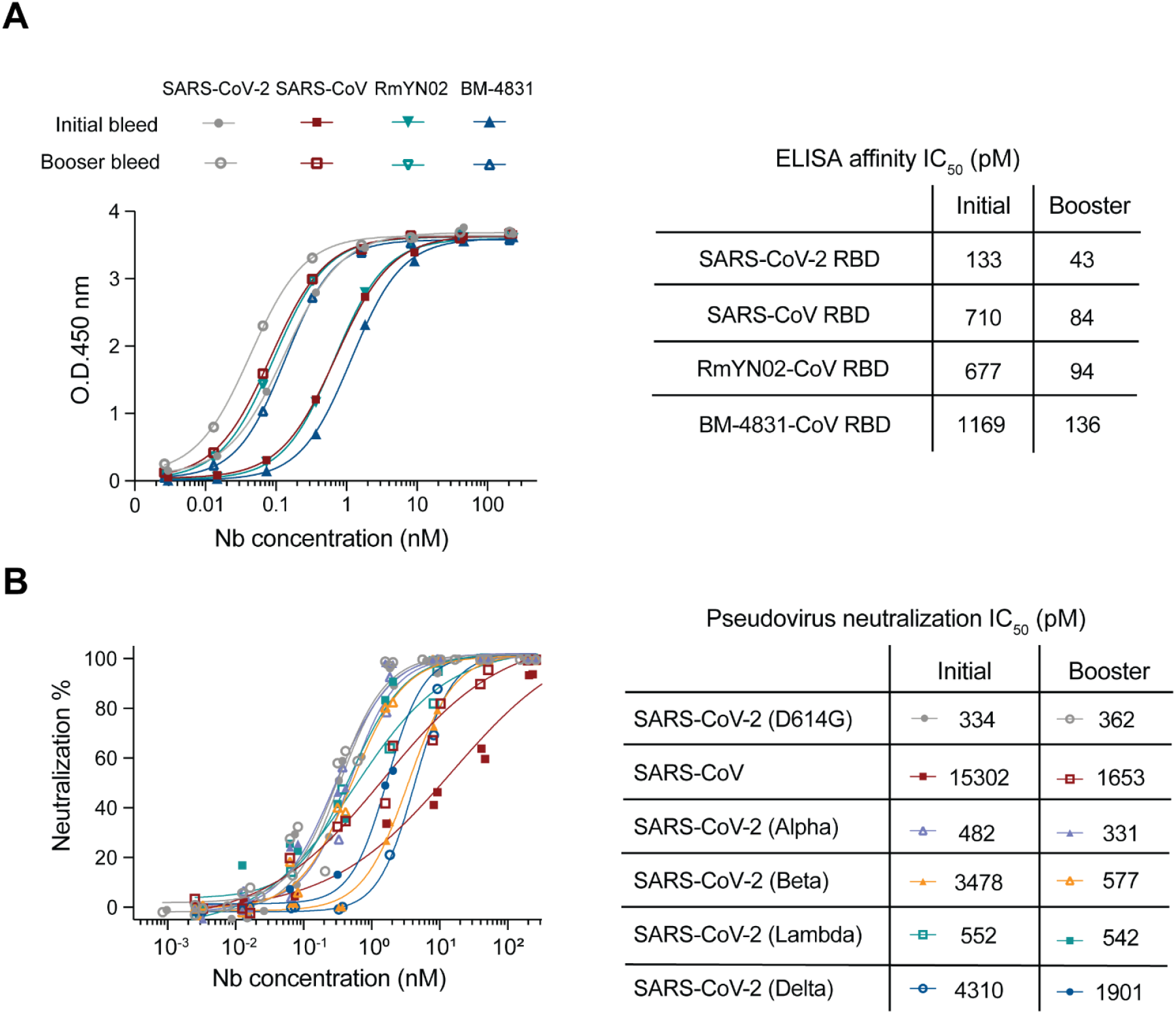
Analysis of the total V_H_Hs isolated from serum after RBD immunization. (**A**) ELISA binding of total V_H_Hs from the initial and booster bleeds against four representative RBDs. (**B**) Pseudovirus neutralization assay of total V_H_Hs of the initial and booster bleeds against SARS-CoV-2, SARS-CoV-2 variants and SARS-CoV.

**Fig. S2.**
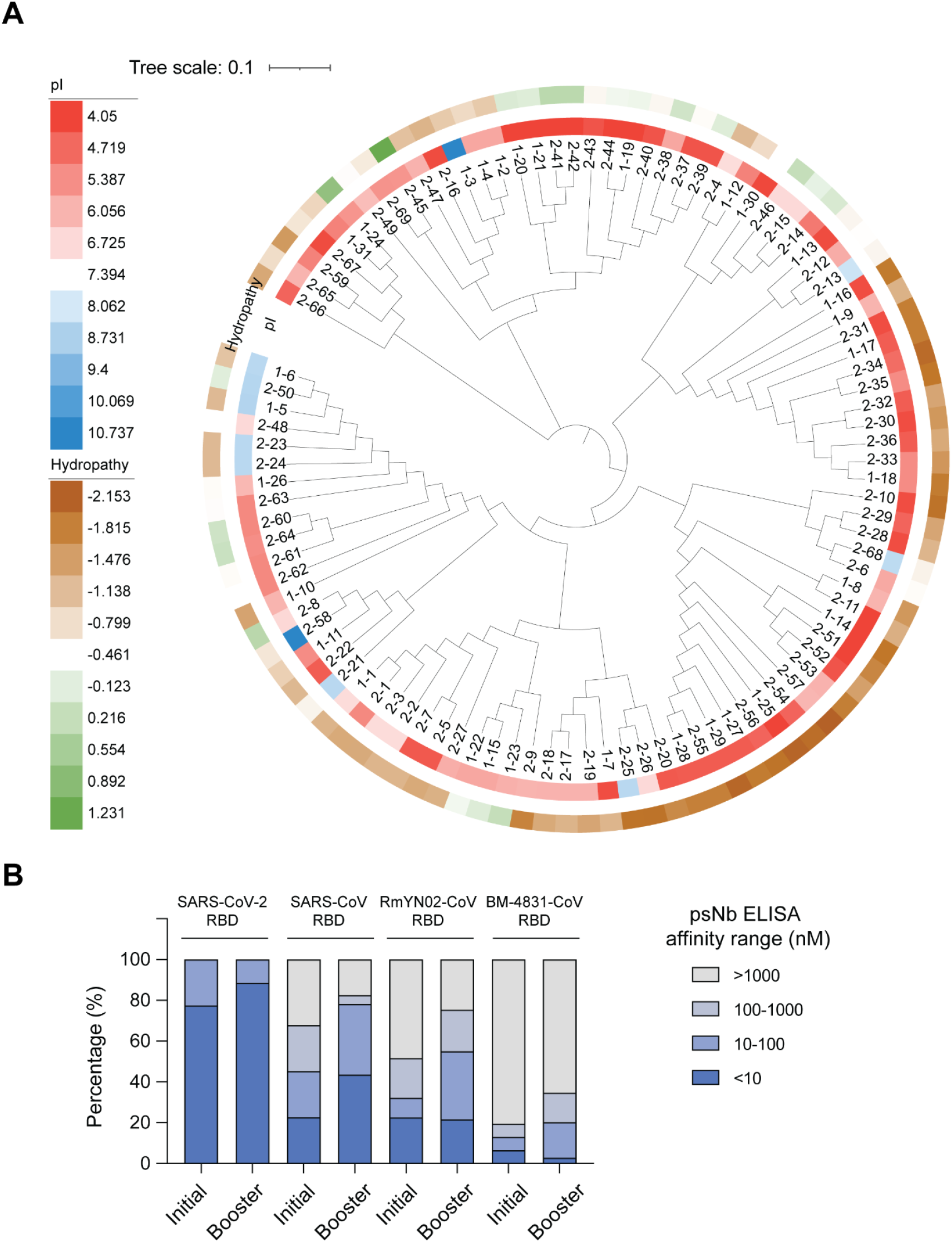
Analysis of experimentally verified psNbs. (**A**) Phylogenetic analysis of the psNb CDR sequences and their physicochemical properties including isoelectric point (pI), hydropathy. (**B**) The individual psNb isolated from either initial or booster bleed was synthesized, purified and evaluated for RBDSARS-CoV-2, RBDSARS-CoV, RBDRmYN02-CoV and RBDBM-4831-CoV binding by ELISA. The percentage was calculated as the number of psNbs in a certain affinity range divided by the total number of the psNbs.

**Fig. S3.**
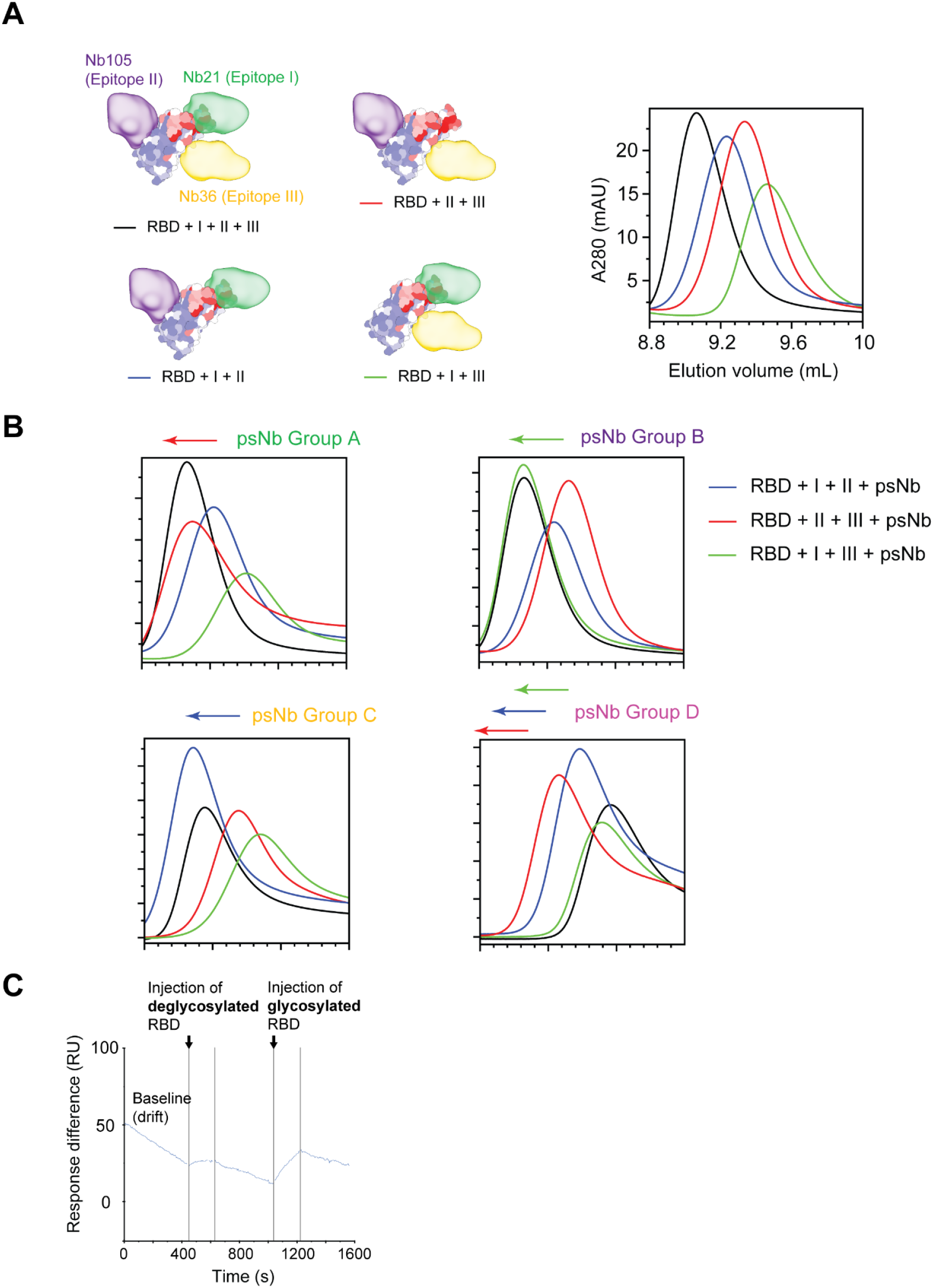
Size exclusion chromatography (SEC) and surface plasmon resonance analysis of representative psNbs for RBD binding (summarized in **table S1**). (**A**) SEC profiles of the reconstituted nanobodies of non-overlapping epitopes in complex with RBD_SARS-CoV-2_. Left panel shows the structures that help illustrate competitive SEC experiments. (**B**) Competitive SEC of psNbs. Group A psNbs compete with Nb21. Group B psNbs compete with Nb105. Group C psNbs compete with Nb36. Group D psNbs do not compete with any of these three Nbs. Group E psNbs dissociate from RBD (**Methods**). (**C**) A representative SEC unclassified psNb (2-47) was covalently coated to the flow cell (Fc) 2 of a CM5 sensorchip. Fc1 of the sensorchip was non-coated as control. 1 µM deglycosylated RBDSARS-CoV-2 in the HBS-EP buffer was injected to the surface at 20 µl/min for 3 mins with dissociation for 3 mins, followed by a sequential injection of 1 µM glycosylated RBDSARS-CoV-2. The responses were recorded as the difference between the signals from Fc2 and Fc1.

**Fig. S4.**
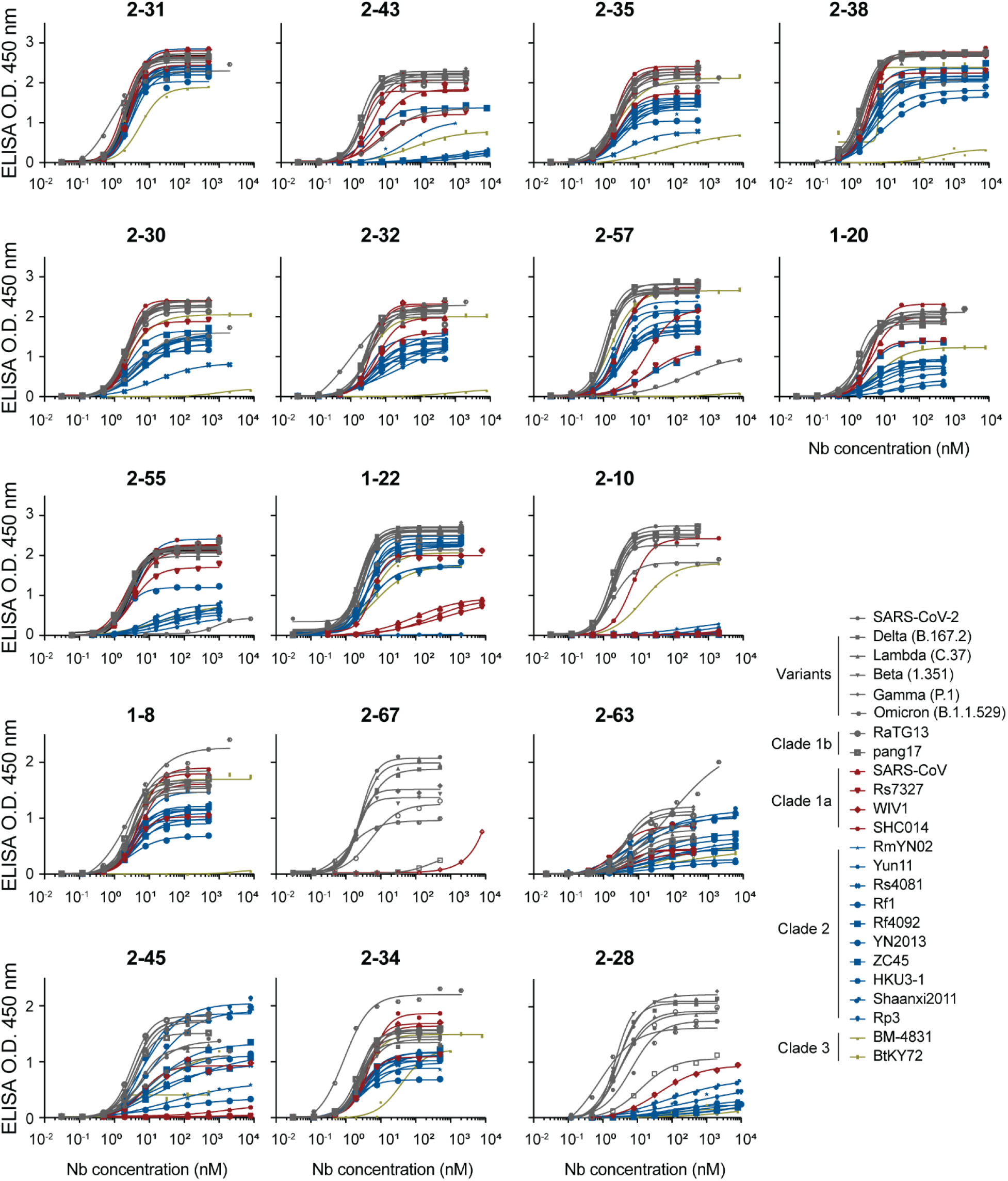
ELISA of 17 psNbs against the RBDs from the WT SARS-CoV-2, variants and 18 different sarbecoviruses that span all four clades.

**Fig. S5.**
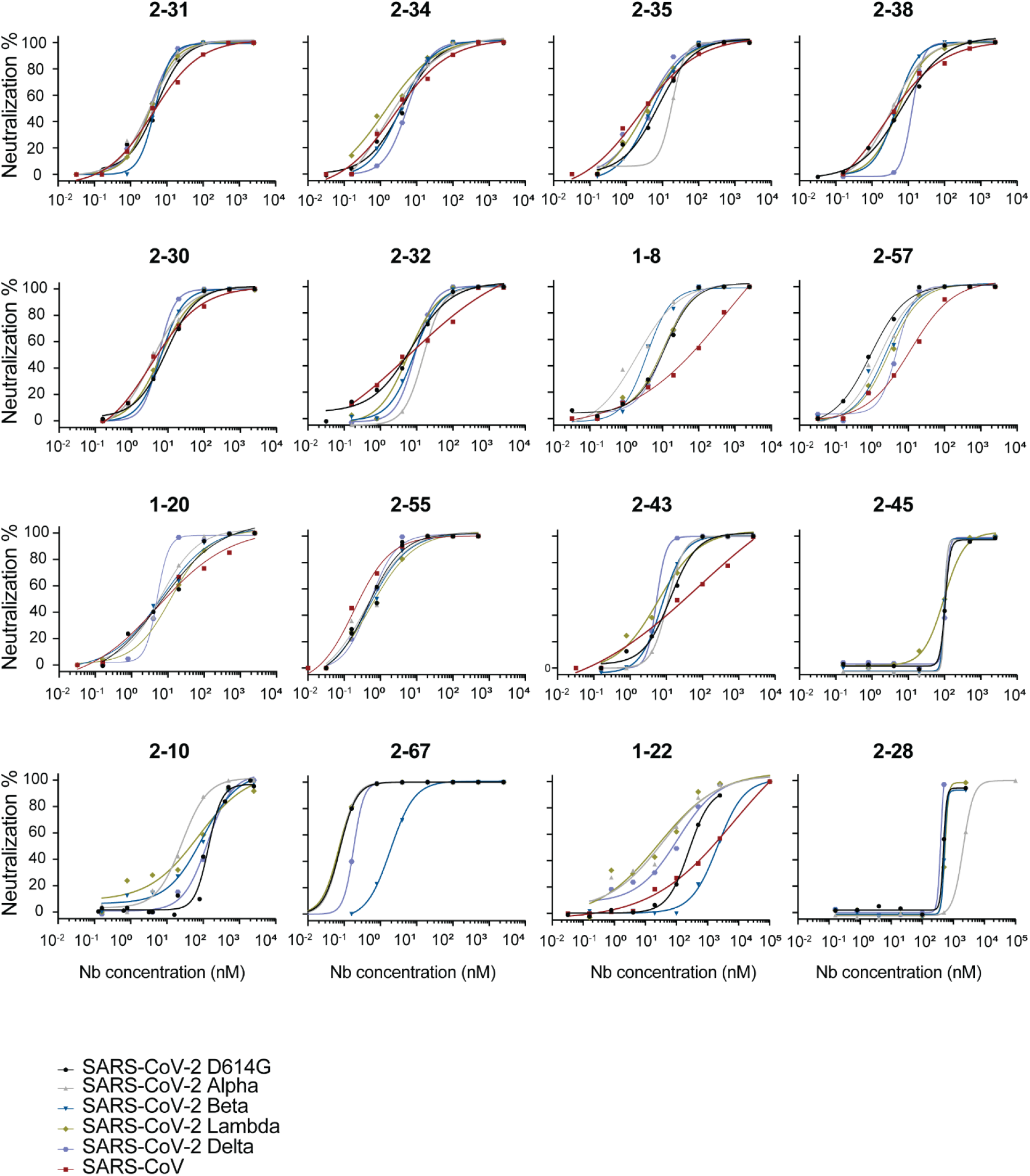
Neutralization assays of 16 psNbs against pseudotyped SARS-CoV-2 (Wuhan-Hu-1, D614G), the VOCs, VOI and SARS-CoV.

**Fig. S6.**
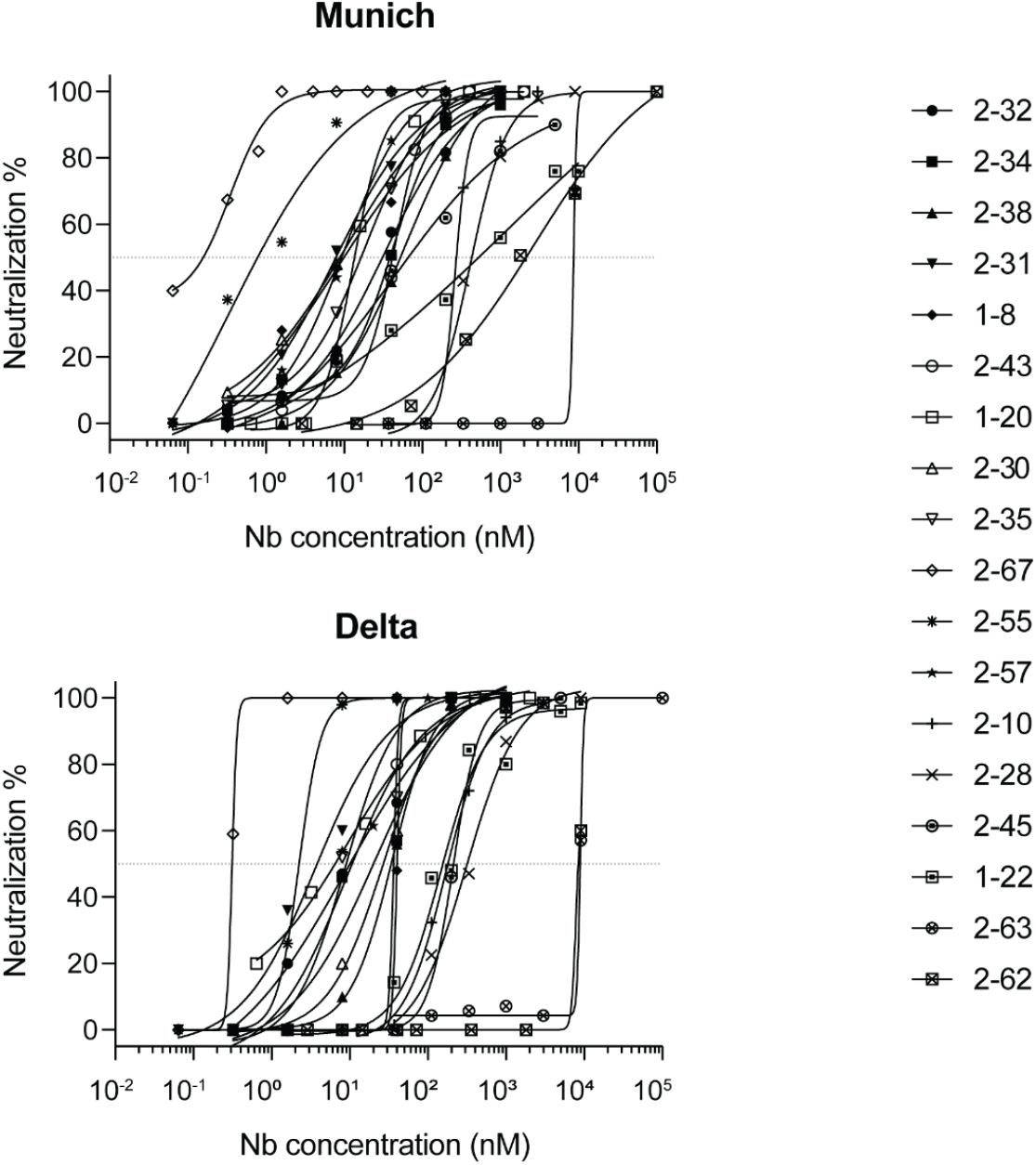
PRNT assay of 17 presentative psNbs against SARS-CoV-2 (Munich strain) and Delta variant.

**Fig. S7.**
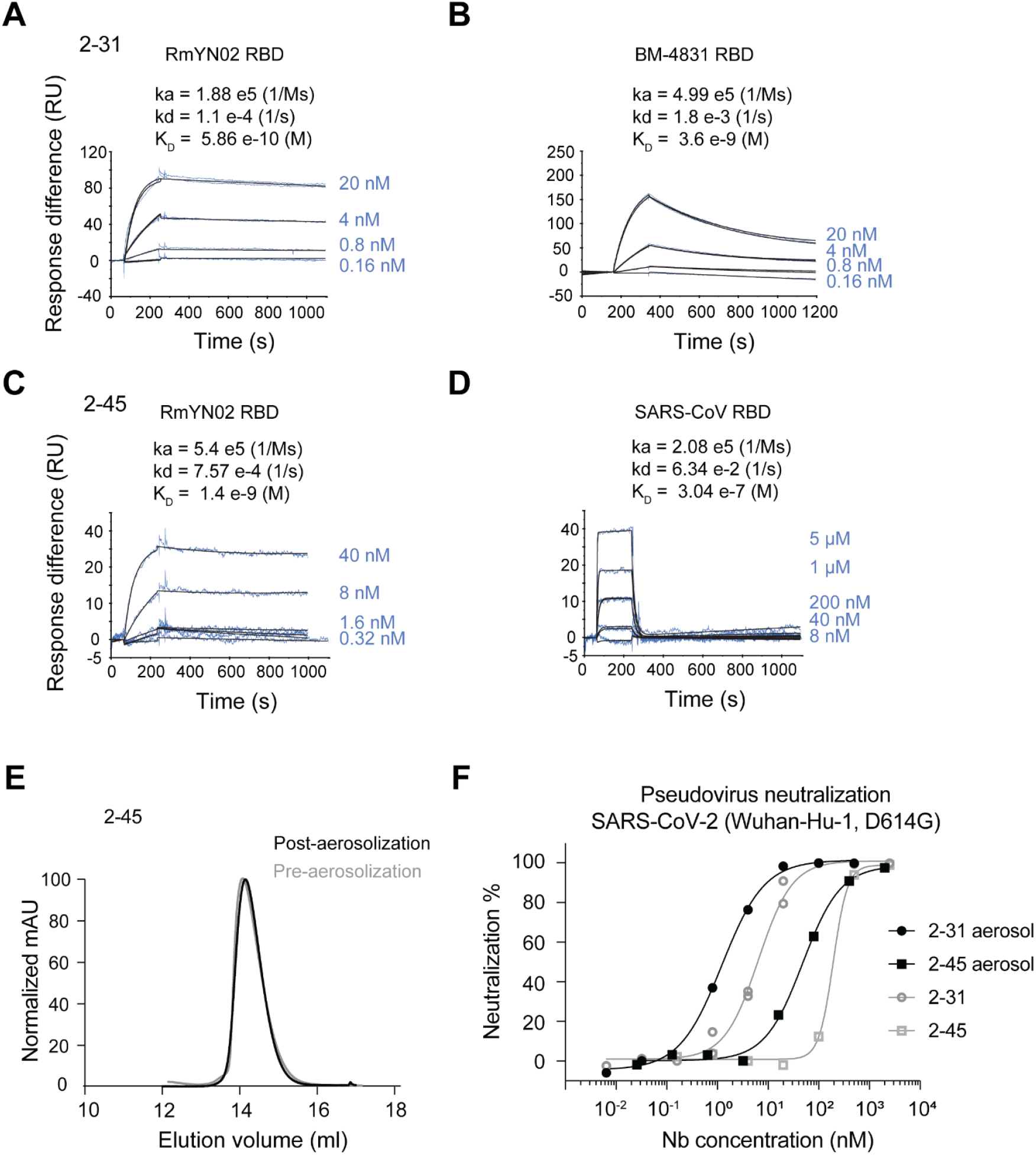
Biophysical characterizations of highly broad psNbs (2-31 and 2-45). (**A-B**) SPR kinetic measurement of 2-31 against RBDRmYN02 and RBDBM-4831. (**C-D**) SPR kinetic measurement of 2-45 against RBDSARS-CoV RBDBM-4831. (**E**) SEC analysis of 2-45 pre and post aerosolization. (**F**) Pseudovirus neutralization of 2-31 and 2-45 pre and post aerosolization.

**Fig. S8.**
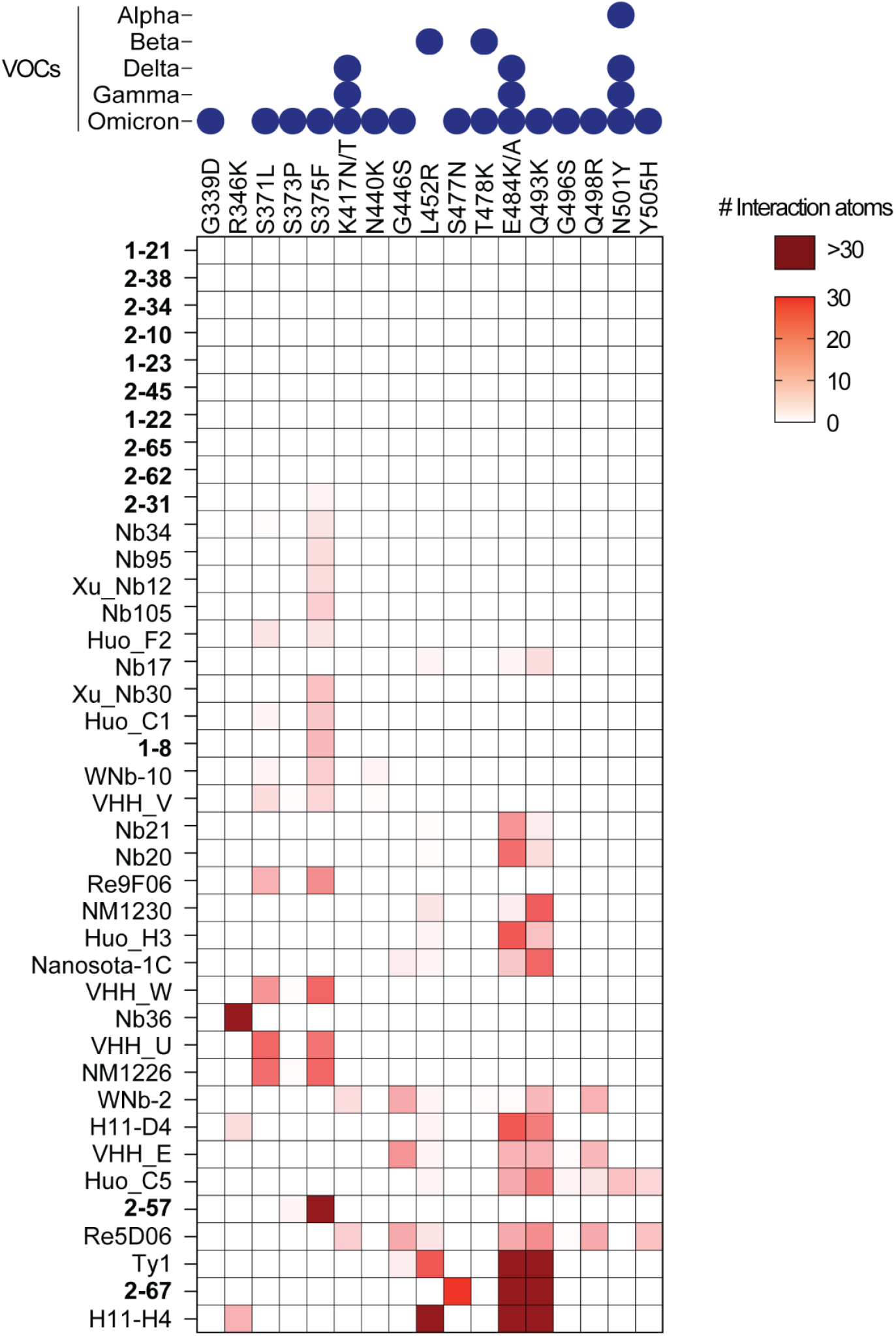
Structural analysis of Nb binding to the RBD mutations from SARS-CoV-2 VOCs. Interfaces were extracted from PDB Nb structures. The number of Nb-interacting atoms per each VOC residue was counted to generate the plot. The psNbs identified in this study are bolded.

**Fig. S9.**
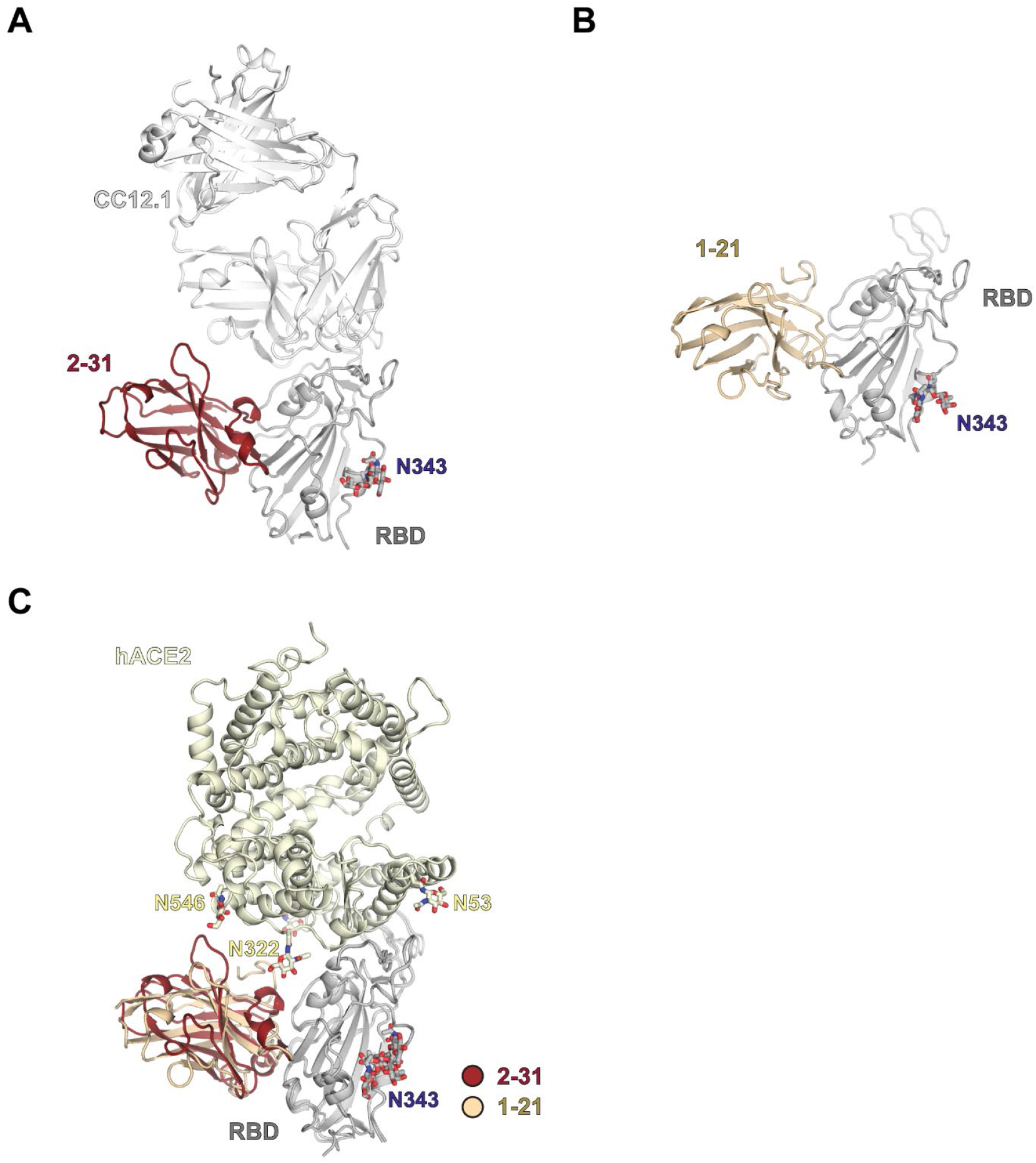
Comparison of overall structures of 2-31 and 1-21 bound to SARS-CoV-2 RBD. Structures are shown as ribbon representation. Glycans in SARS-CoV-2 RBD and ACE2 are shown in sticks and labeled with their corresponding asparagine(N) position number. CC12.1 was used to complex with 2-31 and SARS-CoV-2 to obtain diffraction-quality crystals for structure determination. 2-31 is shown in dark red, 1-21 in beige, RBD in light gray, ACE2 in light yellow, and CC12.1 in white. Structures of 2-31, 1-21, and ACE2 bound to SARS-CoV-2 are superimposed for comparison in **c**.

**Fig. S10.**
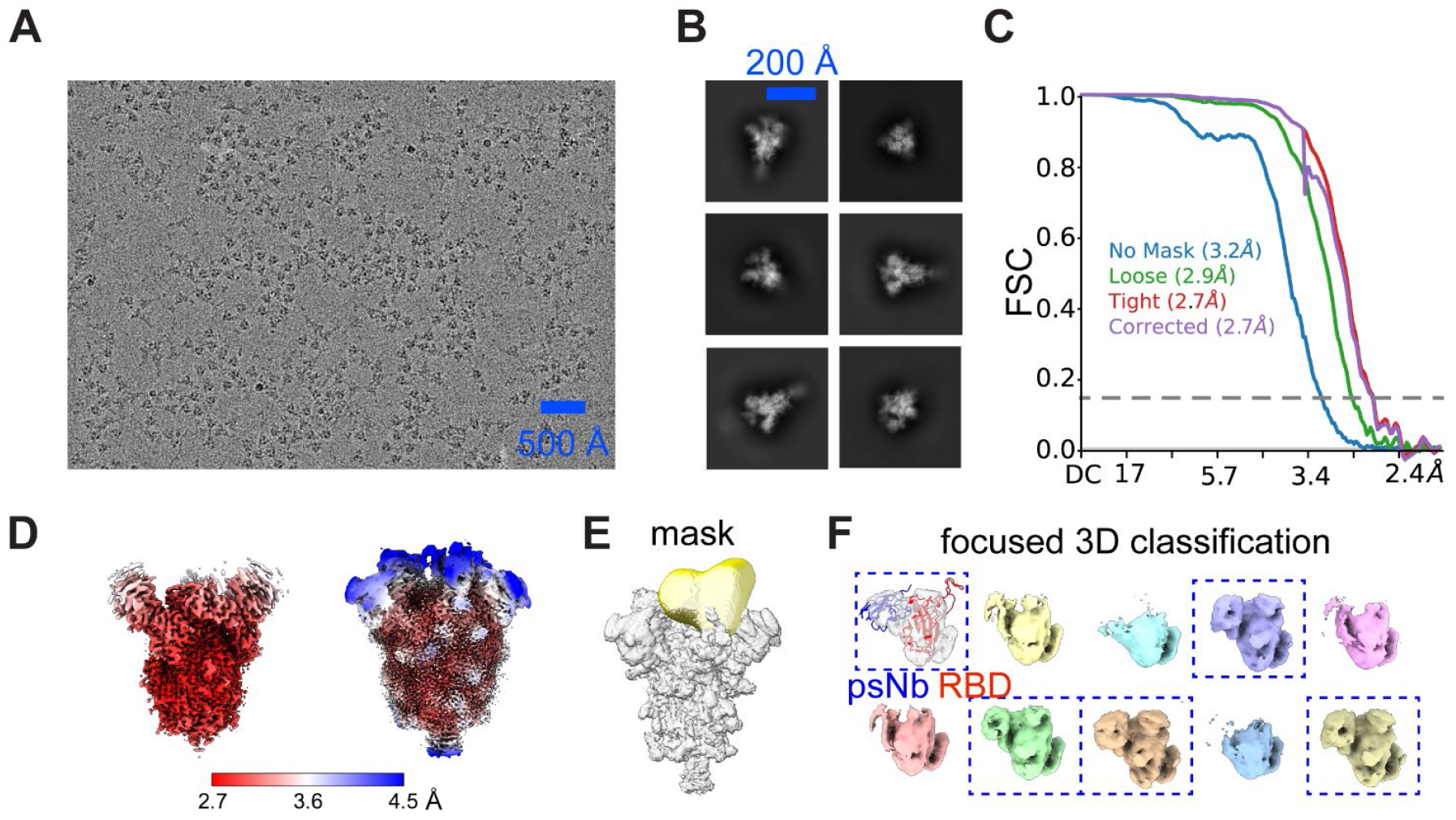
Cryo-EM Structure determination workflow illustrated by 2-57. (**A**) Representative micrograph of spike with 2-57. (B) Representative 2D class averages of the spike glycoprotein with 2-57. (**C**) Gold-standard Fourier shell correlation (FSC) curves calculated from two independently refined half-maps with different masks for spike with 2-57. (**D**) Local resolution map shown at two density thresholds (left: 0.5 and right: 0.1). (**E**) Masking of the RBD:psNb density for focused classification. (**F**) Results of focused classification. Classes with strong RBD:psNb densities (blue rectangle boxes) are selected for further local refinements.

**Fig. S11.**
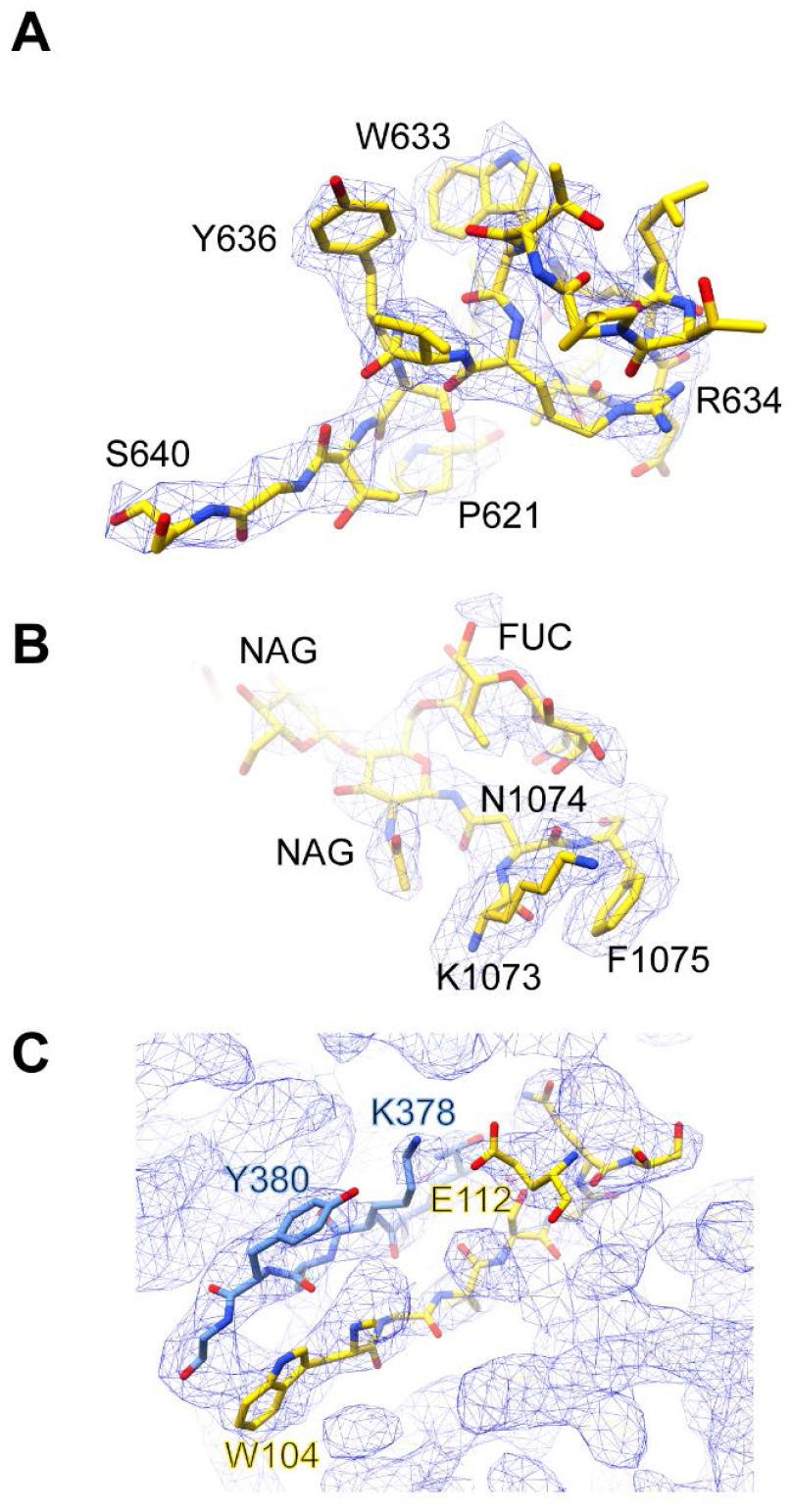
Representative cryo-EM densities of psNbs in complex with the spike or RBD. (**A**) High-resolution cryo-EM maps allow the modeling of residue 621 to 640 for the spike protein, which are absent in the starting structure (PDB: 7CAK). The map was contoured at level 0.15. (**B**) High-resolution cryoEM maps enable the delineation of complex glycan composition at Asn1074 with a Fucose (FUC) attached to the first N-acetylglucosamine (NAG). The map was contoured at level 0.15. (**C**) High-resolution cryoEM maps facilitated modeling of the RBD (blue): 2-57 (yellow) interface. The map was contoured at level 0.64.

**Fig. S12.**
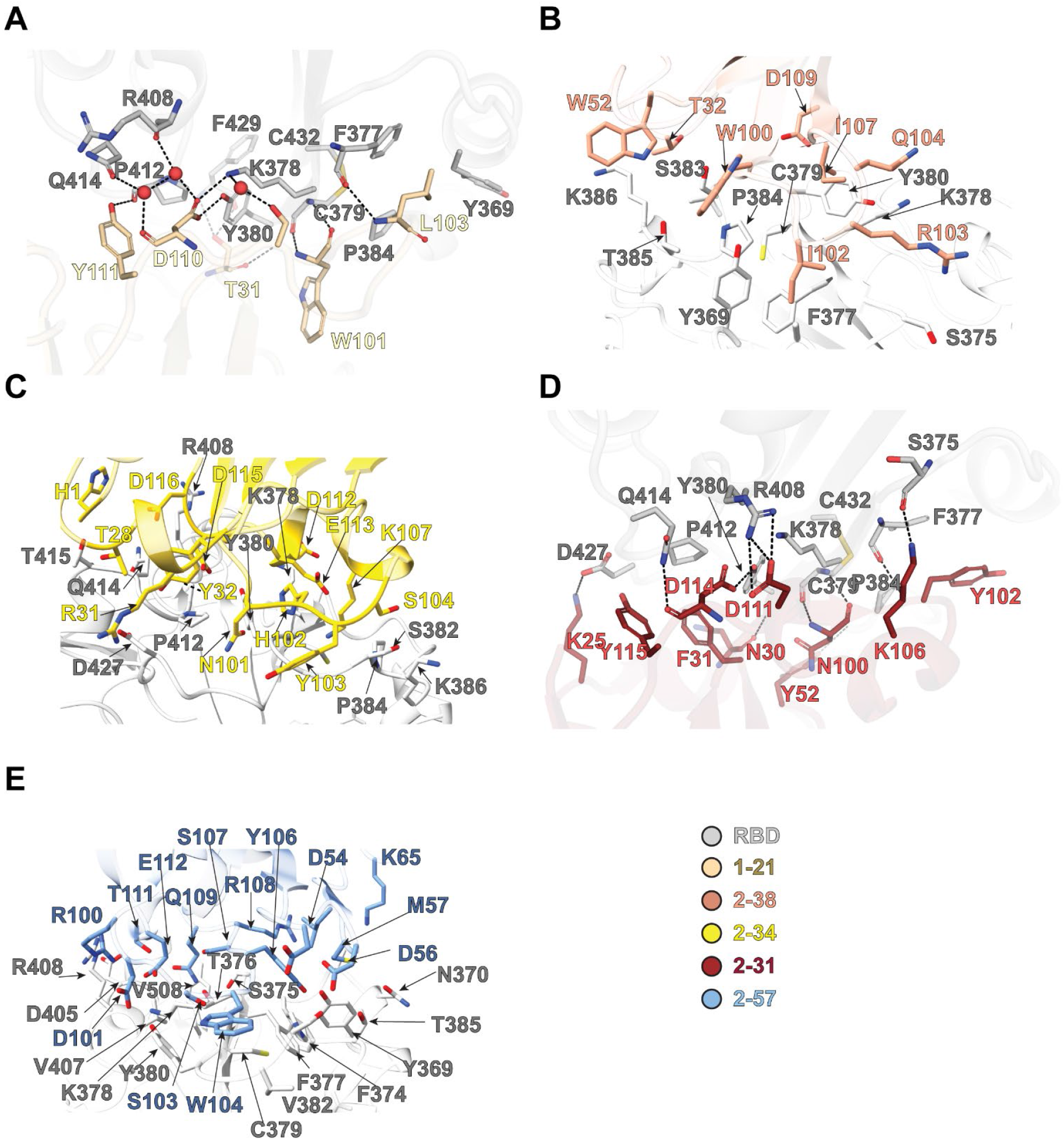
Interactions of epitope II psNbs in complex with the spike or RBD. Key residues involved in the RBD: psNb interfaces are shown in sticks. Hydrogen bonds or salt bridges are indicated with dashed lines. Water molecules mediating interaction between 1-21 and SARS-CoV-2 RBD are shown in red spheres. 1-21 in navajo white, 2-38 in dark salmon, 2-34 in gold, 2-31 in reddish brown, 2-57 in cornflower blue and RBD in light gray. (**A**) Interactions between RBD and 1-21. (**B**) Interactions between RBD and 2-38. (**C**) Interactions between RBD and 2-34. (**D**) Interactions between RBD and 2-31. (**E**) Interactions between RBD and 2-57.

**Fig. S13.**
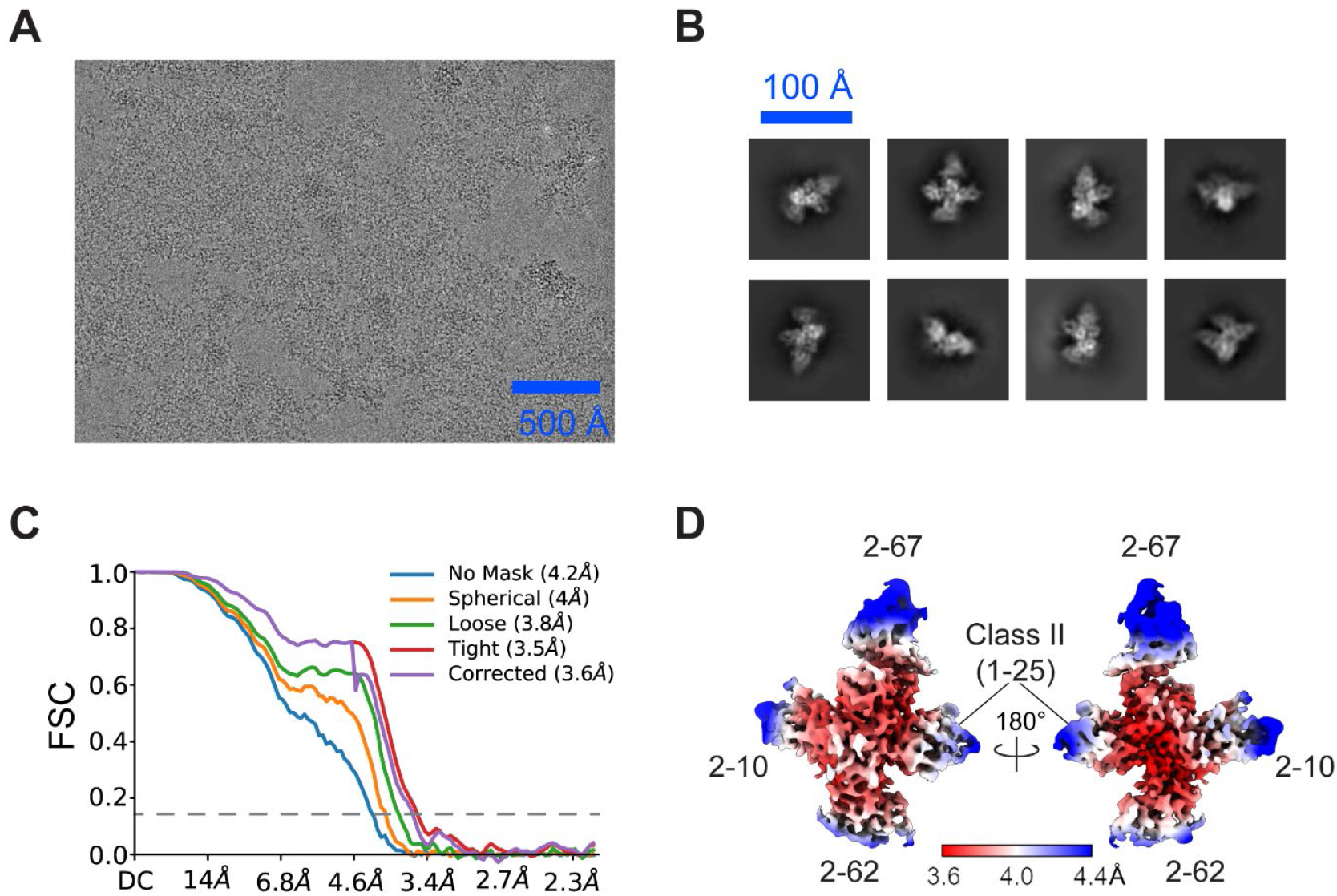
Cryo-EM Structure determination of RBD with 2-67, 2-10, 2-62 and a class II psNb (1-25) and focused refinement. (**A**) Representative micrograph of RBD with four psNbs. (**B**) Representative 2D class averages of RBD and psNbs complex. (**C**) Gold-standard Fourier shell correlation (FSC) curves calculated from two independently refined half-maps with different masks for RBD and psNbs complex. (**D**) Local resolution map contoured at level 0.36.

**Fig. S14.**
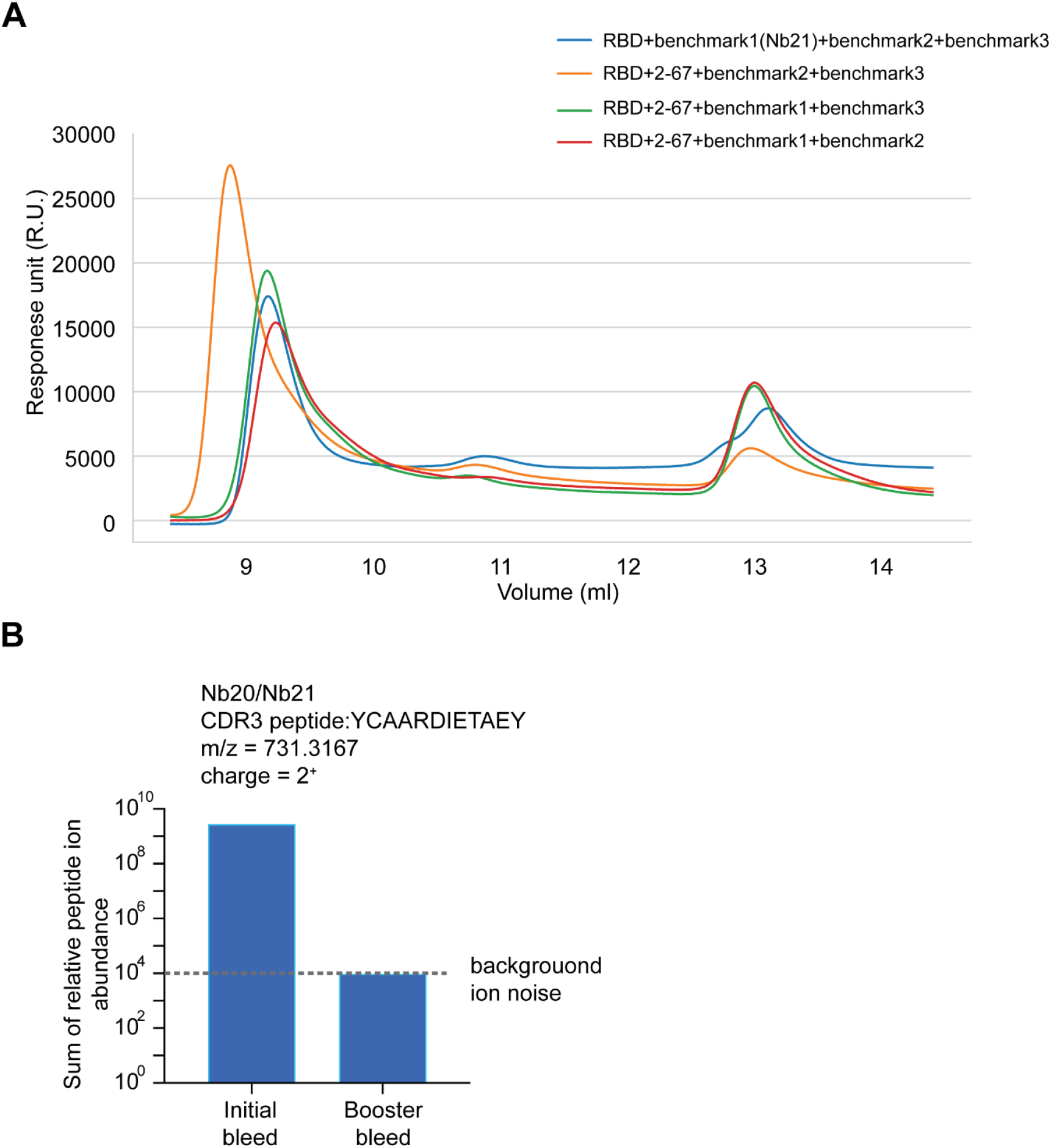
SEC analysis of 2-67 with RBD and benchmark Nbs, and quantitative MS evaluation of Nbs 20 and 21. (**A**) Competitive SEC analysis of 2-67 with Nb21 and other high-affinity benchmark Nbs for RBD binding. (**B**) MS quantification of a CDR3 tryptic peptide specific to Nbs 20 and 21. The Y axis represents the total MS1 ion abundance (relative abundance) of the peptide. Signals below 10^4-5^ are generally considered chemical backgrounds.

**Fig. S15.**
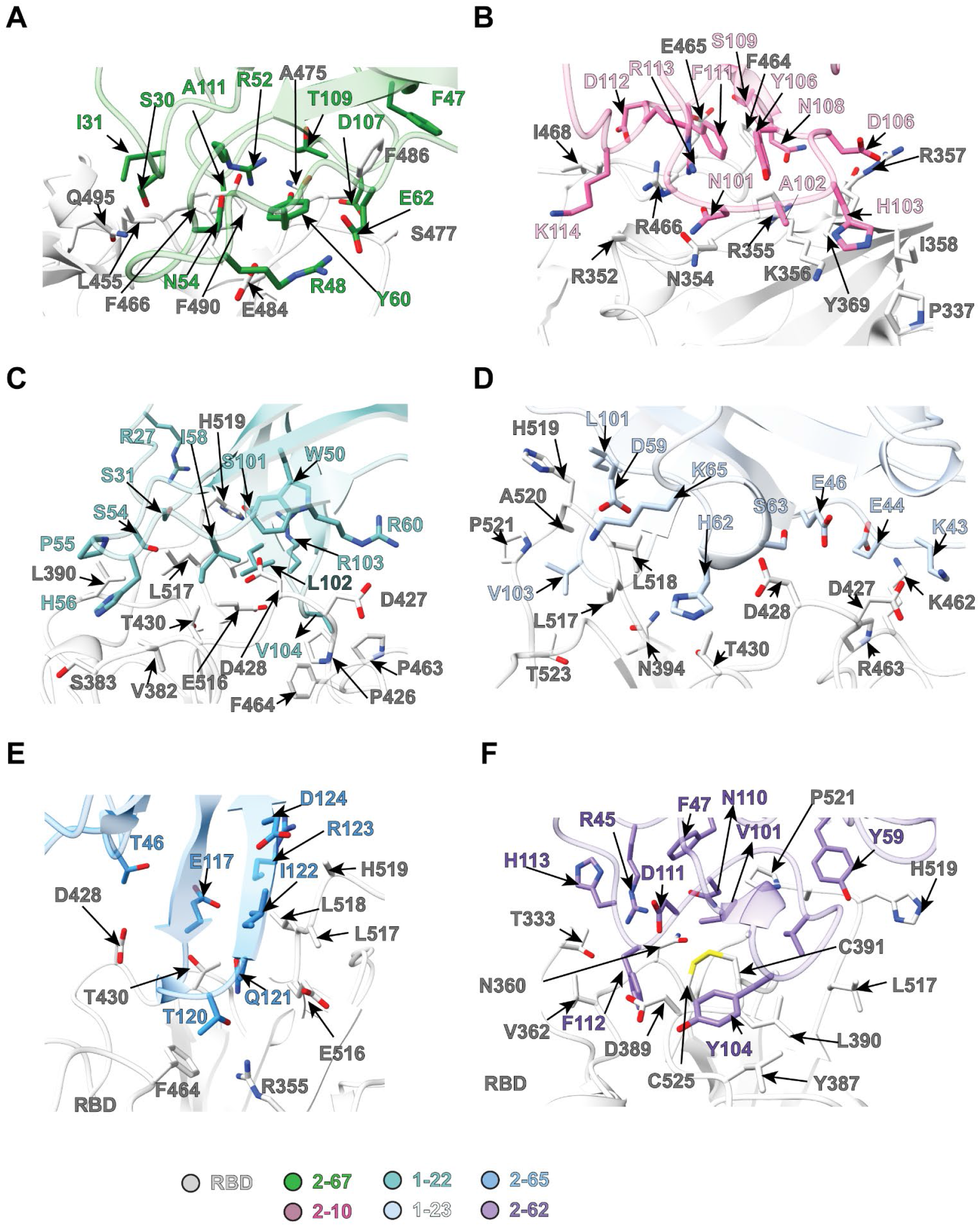
Interactions of class I, III, IV and V psNbs in complex with SARS-CoV-2 spike or RBD. (**A**) Interactions between RBD and 2-67. (**B**) Interactions between RBD and 2-10. (**C**) Interactions between RBD and 1-22. (**D**) Interactions between RBD and 1-23. (**E**) Interactions between RBD and 2-65. (**F**) Interactions between RBD and 2-62.

**Fig. S16.**
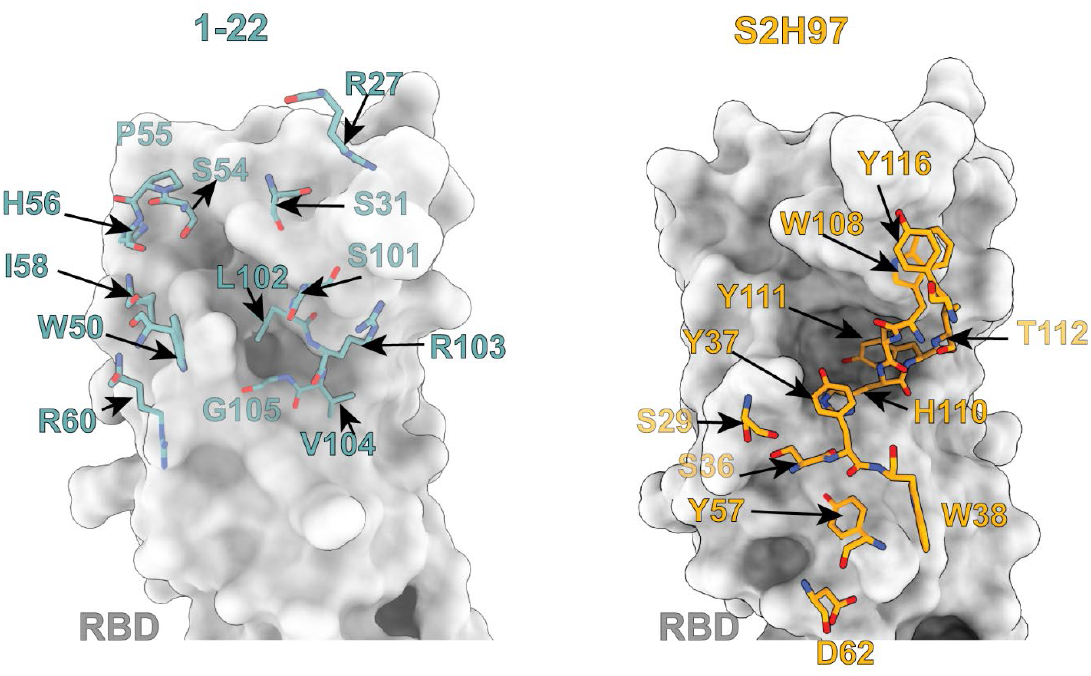
Structural comparison of 1-22 and an IgG antibody (S2H97) binding to the RBD_SARS-CoV-2_.

**Fig. S17.**
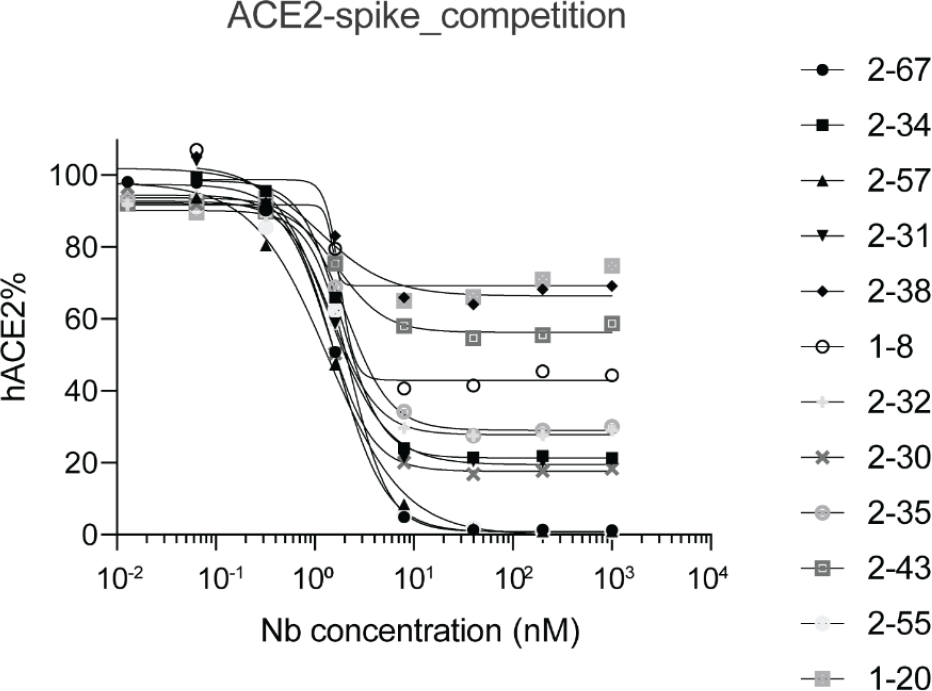
hACE2 competition assay with SARS-CoV-2 spike by ELISA.

**Fig. S18.**
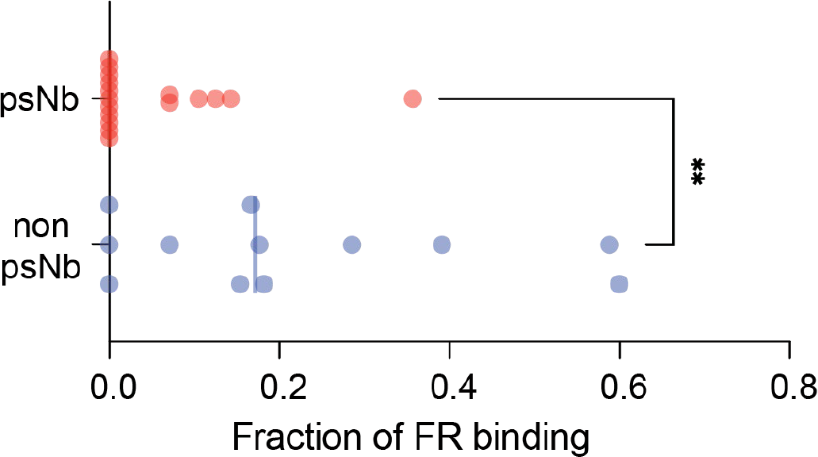
Comparison of the utility of framework residues of psNbs and non-psNbs for binding to RBD.

**Fig. S19.**
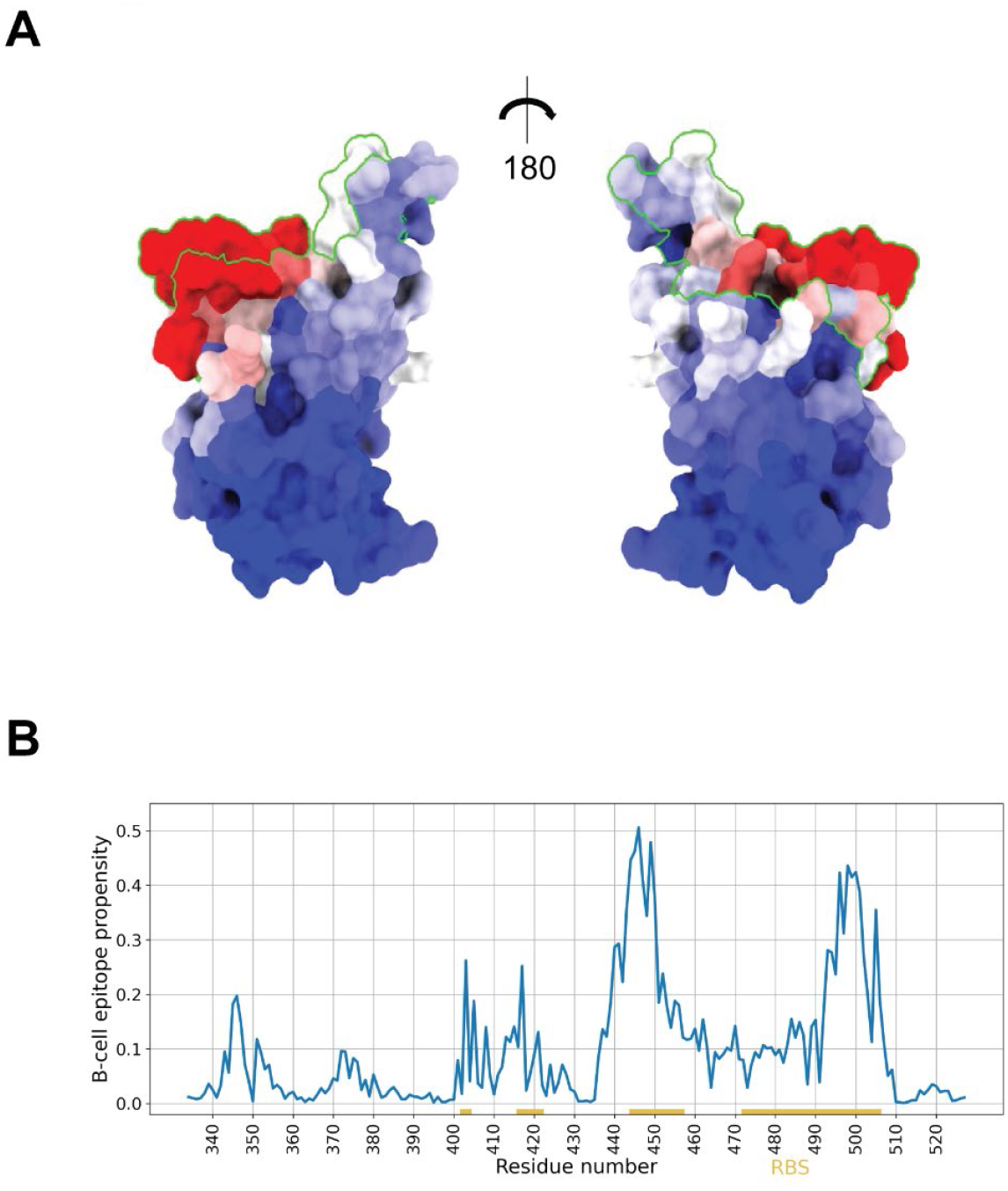
RBD epitope prediction by ScanNet. (**A**) The RBD surface is colored by epitope propensity (from blue = 0.0 to red = 0.35 and higher) using ChimeraX. The ACE2 binding site is traced by a green line. (**B**) RBD epitope propensity profile.

**Fig. S20.**
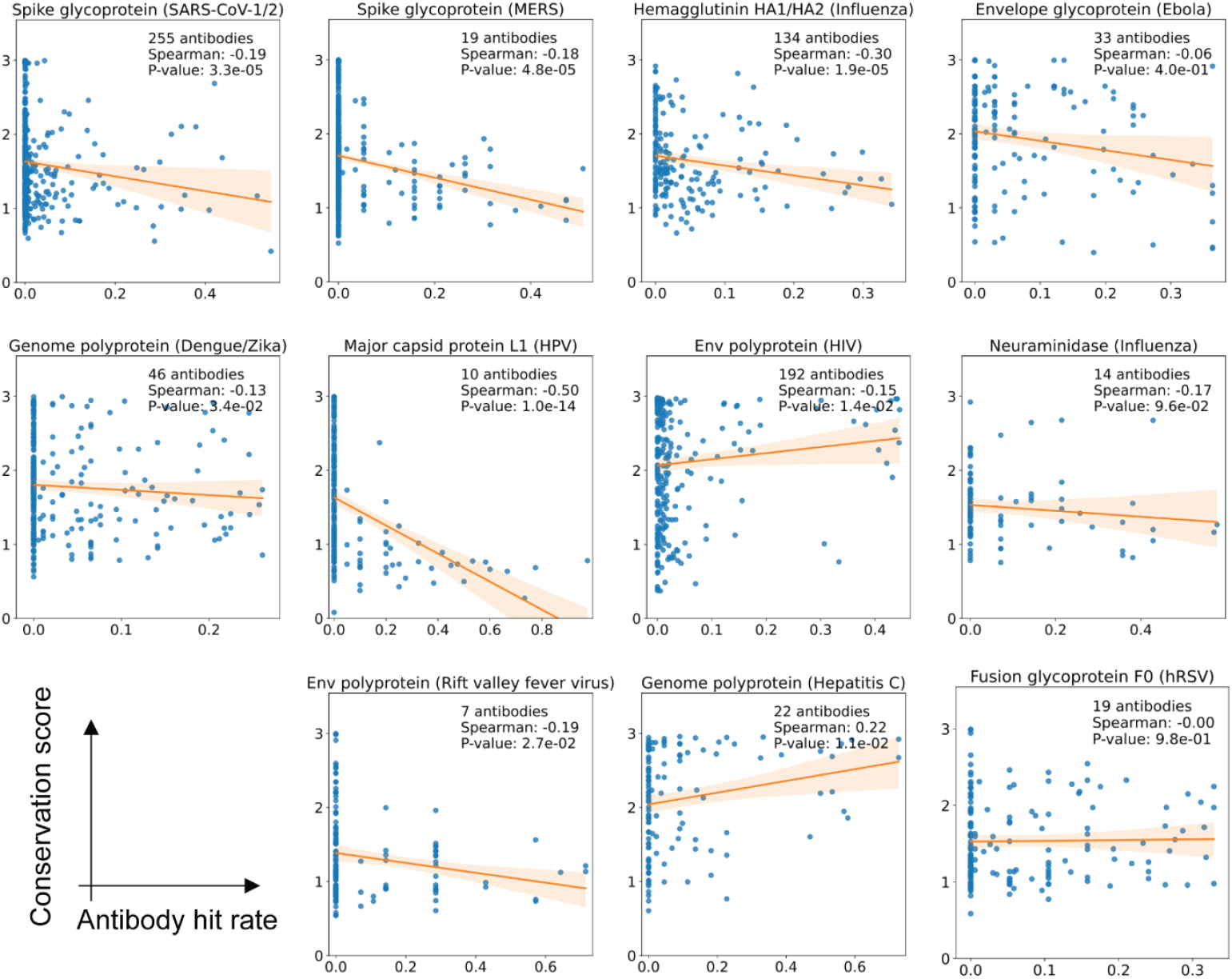
Scatter plots of antibody hit rates against conservation scores for surface residues for 11 antigens. A negative trend is found for most antigens. Notable outliers include the envelope protein of HIV and the genome polyprotein of Hepatitis C; we speculate that this is due to (i) over-representation of broadly neutralizing antibodies in the PDB and/or (ii) extensive glycosylation.

**Fig. S21.**
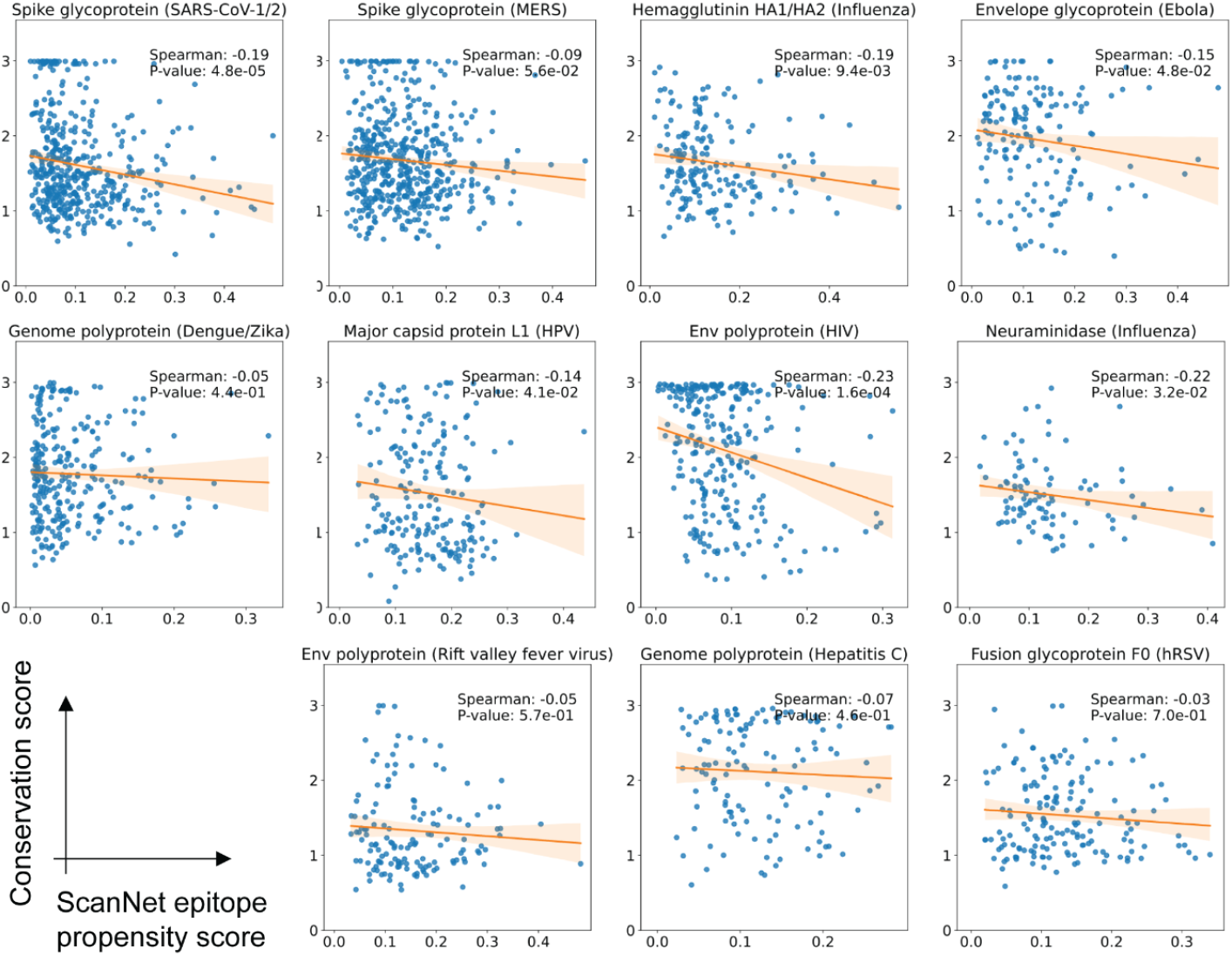
Scatter plots of ScanNet epitope propensity scores against conservation scores of surface residues for 11 antigens. A negative trend is observed between the conservation score and the ScanNet epitope propensity score which was calculated from antigen structure only. This suggests that the association between conservation and antibody hit rate arises solely from structural properties.

